# A Saposin deficiency model in *Drosophila*: lysosomal storage, progressive neurodegeneration, sensory physiological decline and defective calcium homeostasis

**DOI:** 10.1101/078725

**Authors:** Samantha J. Hindle, Sarita Hebbar, Dominik Schwudke, Christopher J. Elliott, Sean T. Sweeney

**Author notes:** **Correspondence:** Sean T. Sweeney, Department of Biology, University of York, Wentworth Way, Heslington, York, YO10 5DD, U.K. Present address: Department of Anesthesia and Perioperative Care, Genentech Hall, 600 16^th^ Street, University of California San Francisco, San Francisco, CA. Max Planck Institute of Cell Biology and Genetics, Dresden, 01307, Germany (Sarita Hebbar). Research Center Borstel, Leibniz-Center for Medicine and Biosciences, Borstel, 23845, Germany (Dominik Schwudke).

## Abstract

Saposin deficiency is a childhood neurodegenerative lysosomal storage disorder (LSD) that can cause premature death within three months of life. Saposins are activator proteins that promote the function of lysosomal hydrolases in the degradation of sphingolipids. There are four saposin proteins in humans, which are encoded by the *prosaposin* gene. Mutations causing an absence of individual saposins or the whole prosaposin gene lead to distinct LSDs due to the storage of different classes of sphingolipids. The pathological events leading to neuronal dysfunction induced by lysosomal storage of sphingolipids are as yet poorly defined. We have generated and characterised a *Drosophila* model of saposin deficiency that shows striking similarities to the human diseases. *Drosophila saposin-related* (*dSap-r*) mutants show a reduced longevity, progressive neurodegeneration, lysosomal storage, dramatic swelling of neuronal soma, perturbations in sphingolipid catabolism, and sensory physiological deterioration. We have also revealed a genetic interaction with a calcium exchanger (CalX), suggesting that calcium homeostasis may be altered in saposin deficiency. Together these findings support the use of *dSap-r* mutants in advancing our understanding of the cellular pathology implicated in saposin deficiency and related LSDs.

## Introduction

Saposin deficiency is an autosomal recessive lysosomal storage disorder (LSD) that is typically associated with severe, age-dependent neurodegeneration and premature death during early childhood. In humans there are four saposins (saposins A – D), which are encoded by the *prosaposin* gene (1, 2, 3). The mature, active saposins are produced by cleavage of the prosaposin precursor during passage through the endosomes to the lysosomes; this function is primarily performed by Cathepsin D (4). Once in the acidic lysosome environment, saposins act as activator proteins and promote the function of hydrolases involved in sphingolipid degradation (5-9). Mutations in *prosaposin* therefore cause a primary accumulation of sphingolipid species in the lysosomes. The location and severity of the *prosaposin* mutation dictates the number of saposins that are affected and hence the degree of sphingolipid accumulation and age of lethality. Mutations abolishing the *prosaposin* start codon result in an absence of prosaposin and therefore all 4 saposins; this causes the most severe pathology and individuals present with severe neurodegeneration at birth and die within 4 months (10-12). Of the single saposin disorders, saposin A deficiency is the most severe and results in death at 8 months old (13), whereas the mildest of the saposin C mutations cause non-neuronopathic disorders with relatively mild symptoms until the fourth decade of life (14). No single saposin D deficiencies have been reported in humans.

Because each saposin generally promotes the function of a specific sphingolipid hydrolase, the single saposin deficiencies resemble the pathology caused by mutations in their cognate hydrolase (e.g. 15; reviewed in 16); total prosaposin deficiency encapsulates many aspects of the single saposin deficiencies but to a more severe degree (10-12).

The sphingolipdoses form the largest group of LSDs, yet to date only one sphingolipidosis (Niemann-Pick Type C (NPC)) has been modelled in *Drosophila* (17-20). To broaden our understanding of the sphingolipidoses, and to help identify pathological events subsequent to sphingolipid storage, we generated a *Drosophila* model of saposin deficiency. The *Drosophila Saposin-related* (*dSap-r*) locus encodes a protein predicted to contain eight saposin-like domains, each containing the classic six-cysteine arrangement found in all mammalian saposins. Characterisation of *dSap-r* mutants revealed pathology similar to that of the human disorders, including reduced longevity, progressive neurodegeneration, aberrant sphingolipid levels, and physiological deterioration; all hallmark signs of lysosomal storage. Our analysis reveals a genetic interaction with the Na+/Ca+ exchanger, CalX, and suggest a deficit in calcium regulation in the *Drosophila* model of saposin deficiency.

## Materials and Methods

### Identification of *Drosophila Sap-r*

A blastp search (NCBI; www.ncbi.nlm.nih.gov) was performed to identify the *Drosophila melanogaster* prosaposin (PSAP) homologue. The entire *Homo sapiens* PSAP protein sequence (CAG33027) was used to search the *D. melanogaster* protein database. Standard blastp assumptions were applied. A reciprocal search was performed, using the *D. melanogaster* d-Sap-rPA sequence to blast the *H. sapiens* protein database, to ensure the correct homologue was identified.

### Identification of *Drosophila Sap-r* monosaposins

To identify the putative monosaposins encoded by the *dSap-r* gene, each human monosaposin sequence (RCSB Protein Data Bank entries 2DOB, 1N69, 2GTG, 2RB3) was aligned against the full-length dSap-rPA sequence using the bl2seq tool at NCBI. The *H. sapiens* and *D. melanogaster* monosaposins were aligned using default settings in ClustalX.

#### *Drosophila* stocks

All experimental crosses were grown on maize-meal fly food at 25°C. Newly eclosed flies were transferred to standard yeast-sucrose-agar fly food. The wild type (+/+) control for all experiments was *w^1118^* crossed to Canton-S. The *dSap-r*^*PBac*^ and Df(3R)*tll-e* (subsequently referred to as Df) stocks were from the Bloomington stock centre and the *dSap-r*^*NP7456*^ stock was from the Kyoto stock centre. The *dSap-r*^*C27*^ deletion was generated by mobilising the *P*-element from the *dSap-r*^*NP7456*^ parent line. The *UAS-dSap-r* transgenic stock was generated for this study. Briefly, pUAST-*dSap-r* was generated by excision of *dSap-r* cDNA from the pOT2 vector (clone GH08312, BDGP Gold collection), using XhoI and EcoRI followed by ligation into the pUAST vector. pUAST-*dSap-r* was microinjected into *w^1118^* embryos with helper plasmid *∆2-3*. *UAS-mCD8GFP* and 1407-GAL4 stocks were kindly provided by Andreas Prokop (University of Manchester, UK).

#### Longevity

Newly eclosed flies were collected in separate-sex vials of approximately 10 flies/vial and aged at 29°C (n>100 flies, unless otherwise stated). Flies were transferred to fresh food every 2-3 days and the number of surviving flies recorded. Longevity was plotted as the percentage of Day 0 flies alive each subsequent day.

#### Immunohistochemistry

For the *dSap-r* expression pattern, third instar larvae (n>5) were dissected and fixed in 3.7% formaldehyde/PBS for 10 min, followed by 3 x 5 min washes in 0.1% PBST (PBS with 0.1% Triton X-100). Larvae were labelled overnight at 4°C with mouse anti-repo-8D12 or mouse anti-elav-9F8A9 diluted in PBST (1:50; Developmental Studies Hybridoma Bank, University of Iowa). Washes were performed as above, followed by incubation for 2 h at RT in Cy3-conjugated goat anti-mouse IgG (1:200; Jackson ImmunoResearch). Larvae were washed (as above) and left in 70% glycerol/PBS for 1 – 2 h before mounting in Vectashield (Vector Laboratories). To label lysosomes, aged adult brains were dissected in 4% paraformaldehyde/PBS and transferred to fresh fixative for 20 min (n>8). Brains were washed 3 x 15 - 20 min in 0.3% PBST followed by incubation overnight at RT with rabbit anti-Arl-8 (1:500; kindly provided by Debbie Smith, University of York, U.K) and mouse anti-elav (1:50). Brains were washed as above, followed by incubation for 3 h at RT in FITC-conjugated goat anti-rabbit IgG and Cy3-conjugated goat anti-mouse IgG (1:200; Jackson ImmunoResearch). Brains were washed and mounted in Vectashield. All images were acquired using a Zeiss LSM 510 meta Axiovert 200M laser scanning confocal microscope.

#### Head sectioning

Aged flies were briefly dipped in 30% ethanol before being submerged in fixative (4% paraformaldehyde, 1% glutaraldehyde in 0.1M sodium phosphate buffer, pH 7.4) and pinned through the abdomen. The proboscis and accessible air sacs were rapidly removed from the heads. Heads were transferred to glass vials containing fresh fixative and were vacuum treated to remove trapped/adherent air. Vacuum-treated heads were incubated in fresh fixative overnight at 4°C. All incubations were performed on a rotating wheel, unless otherwise stated. Heads were washed 3 x 10 min in 0.1M sodium phosphate buffer and post-fixed in 1% OsO_4_ for 1 h. Following washes in 0.1M sodium phosphate buffer (3 x 10 min) and dH_2_O (3 x 10 min), heads were dehydrated in an acetone series (30%, 50%, 70%, 90%, 3x 100%; 20 min each). Heads were incubated in increasing concentrations of Spurr’s resin:acetone (25%, 50%, 75%, 95%, 2x 100% [at 37°C]; 45 min each) followed by incubation in 100% resin overnight at 4°C without rotation. Heads were embedded in Spurr’s resin (Spurr, 1969) for 24 h at 70°C. Three semi-thick serial sections (1.0 µm; Leica Ultracut UCT) were taken every 10 – 20 µm until the desired depth was reached. Sections were dried onto glass slides, stained with 0.6% toluidine blue in 0.3% sodium carbonate on a hot plate (80°C) and rinsed with dH_2_O. Sections were imaged using a Zeiss Axiovert 200 microscope equipped with a Zeiss AxioCam HRm camera.

#### Quantification of vacuole number

Vacuoles were quantified manually for the antennal lobes and the visual system (eye, lamina, medulla, lobula and lobula plate) from 3 serial sections per fly head (n≥3). The sections were matched for depth through the head. Vacuoles were counted from both sides of the head. ANOVA statistical tests were performed using SPSS software (IBM Corp., USA).

#### Transmission electron microscopy

After reaching the desired depth by semi-thick sectioning, the same embedded heads used for light microscopy were sectioned for transmission electron microscopy (n=3). Ultrathin sections (60 – 70 nm) were collected on 200 and 400 mesh coated grids, treated with uranyl acetate in 50% ethanol for 10 min and submerged in dH_2_O to wash. Sections were stained with lead citrate for 10 min in the presence of sodium hydroxide pellets, followed by washing in dH_2_O. Images were captured using analysis software on a TECNAI G^2^ (Version 2.18) transmission electron microscope (120 kV).

#### Neuronal soma area quantification

Neuronal soma area was quantified using ImageJ software. The soma and nuclear boundary of each neuron were demarcated and the area calculated by first inputting the number of pixels per micron then using the ImageJ area measurement tool. Neuronal soma area were normalised to nuclear area (n=3 flies). Pseudocolour images were produced using Adobe Illustrator CS4. Student’s t-tests were performed in Microsoft Excel to determine statistical significance.

#### Lipid extraction and lipidomics analyses

Female adults of the required genotypes were collected on emergence and aged for 5 days. Three brains/biological replicate were dissected in PBS, flash-frozen in 20% methanol using liquid nitrogen, and stored at -80°C until lipid extraction was performed. Brain samples were homogenized and extracted according to the methyl-tert-butyl ether extraction method (69). All lipid standards were added to the homogenates prior to extraction (Supplemental Table 2). After phase separation, the organic phase was used for lipidomics and the aqueous phase was processed for determining total protein content using the bicinchoninic acid (BCA) assay (BCA1 kit, Sigma).

For MS measurements, the samples were dissolved in 100 μl methanol containing 0.1% ammonium acetate and subsequently analyzed using a flow-injection system at a flowrate of 1 μl/min and 5 μl sample injection. Negative and positive ion mode spectra were acquired with a LTQ-Orbitrap XL (Thermo Fisher Scientific, Germany) equipped with an Agilent 1200 micro-LC system (Agilent Technologies, USA) and a Nanomate Triversa utilizing 5 μm ID ESI-chips (Advion, Biosciences, USA).

In the negative mode Phosphatidylinositol (PI), Phosphatidylethanolamine (PE), Polyethylene oxide (PE-O), Lysophosphatidylethanolamine (LPE), Phosphatidylcholine (PC), Ceramide phosphorylethanolamine (CerPE), Phosphatidylserine (PS), Phosphatidylglycerol (PG) could be identified according to their accurate mass (70). For PS, the specific neutral loss of 87Da was monitored in the linear ion trap and used for quantification. In the positive mode Sph 14:1, Ceramides and HexCeramides were monitored with MS^3^ in the linear ion trap using the long chain base related fragment ions. All MS^3^ for quantifying sphingolipids were analyzed using Xcalibur software 2.07 (Thermo Fisher Scientific, Germany) while all other analyses were performed using LipidXplorer 1.2.4 (71). Absolute levels of individual lipid species (picomol/ug protein) were summed to arrive at lipid class abundances. A minimum of 4 replicates were used for the analyses. Prism 6 software (Prism Software Corp., USA) was used for graph representation and for determining significant differences applying ANOVA coupled with post-hoc Bonferroni tests.

#### RT-PCR

RNA was extracted from third instar larvae using a QIAGEN RNeasy kit, according to manufacturer’s instructions. RNA was treated with DNase prior to cDNA generation. The following primers were used for RT-PCR: 5’TCCTACCAGCTTCAAGATGAC3’ (rp49 Forward), 5’GTGTATTCCGACCACGTTACA3’ (rp49 Reverse), 5’GCAACTGCAACCTGCTTTCC3’ (dSap-r Forward) and 5’GCATCGTTTCCACCATGTCA3’ (dSap-r Reverse).

All PCR reactions using *rp49* and *dSap-r* primers were performed with an elongation time of 1 min, an annealing temperature of 50°C and 55°C, and a cycle number of 25 and 35, respectively. *dSap-r* primers anneal downstream of the *dSap-r*^*C27*^ deletion, but upstream of the *dSap-r*^*PBac*^ insertion.

#### Western blotting

Soluble protein was extracted from aged flies using 10 µl lysis buffer per fly (150 mM NaCl, 20 mM Tris-HCl, pH 8.0, 2 mM EDTA, 0.5% (v/v) NP-40, 1 complete mini protease inhibitor cocktail (Roche)). A standard western blotting procedure was followed including blocking in 5% milk/TBST (Tris buffered saline supplemented with 0.1% (v/v) Tween-20) followed by incubation in primary then secondary antibody diluted in 5% milk/TBST. Washes were performed using TBST. The following antibodies were used: rabbit anti-Arl-8 (1:2000; kindly provided by Debbie Smith, University of York, UK), mouse anti-β-tubulin E7 (1:100; Developmental Studies Hybdridoma Bank, University of Iowa, USA), mouse anti-cathepsin-L (1:250; R&D Systems), hoseradish peroxidase (HRP)-conjugated goat anti-rabbit (1:6000; Sigma) and HRP-conjugated goat anti-mouse (1:10000; Sigma). Bands were visualised using ECL reagent (GE Healthcare, UK) and developed using a Xograph machine. n=2

#### Behavioural analyses

The climbing ability of female flies was assessed by tapping cohorts of 5 flies to the bottom of a 100 ml glass measuring cylinder and video recording the climbing response over 45 s. The data were analysed by quantifying the climbing speed of each fly over 10% intervals of the measuring cylinder. The maximum speed for each fly was used to calculate the average speed per genotype. The same flies tested at 5-days old were re-tested at 22-days old, when possible (n>20 flies). ANOVA statistical tests were performed using SPSS software (IBM Corp., USA).

#### Electroretinograms

ERGs were performed as described in (72). Briefly, aged female flies were left to climb a trimmed 200 µl pipette tip and were blown to the end leaving the fly head protruding. The fly was fixed into position with nail varnish. Glass electrodes were pulled and filled with *Drosophila* Ringer solution (0.13 M NaCl, 4.7 mM KCl, 1.9 mM CaCl_2_) (34). A recording electrode was placed against one eye and a reference (earth) electrode placed against the proboscis using micromanipulators. The flies were dark-adapted for 5 min (Fig. 6B - C) or 2 min (Fig. 6D - E). ERGs were recorded in response to 5 x 750 ms blue light pulses with 10 s intervals. Light pulses were provided by a blue LED lamp (Kingbright, Taiwan) controlled by DASY*Lab* software (Measurement Computing Corp., USA). ERGs were analysed using DASY*View* software (customised software, C. Elliott) n≥10 flies, unless otherwise stated.

### Supplemental Methods

#### Epifluorescence

Organs were dissected from adult flies in PBS and, in some cases, stained with DAPI. Organs were imaged using a Zeiss stereomicroscope equipped with an Axiocam MRc5 camera, a Neolumar S 1.5x FWD 30 mm lens and a HBO 100 mercury lamp.

## Results

### dSap-r has homologous sequence structure and similar expression pattern to mammalian prosaposin

The *Drosophila* prosaposin (PSAP) homologue (Saposin-related; dSap-r) was identified by a BLAST screen using the complete human PSAP protein sequence (blast E value 4e-36). The *dSap-r* locus is located at band 100A7 of the right arm of the third chromosome. There are two predicted transcripts for *dSap-r* (*dSap-r RA* and *dSap-r RB*); *dSap-r RB* appears to be an in-frame truncation of *dSap-r RA* (Fig. 1A).

**Fig. 1.**
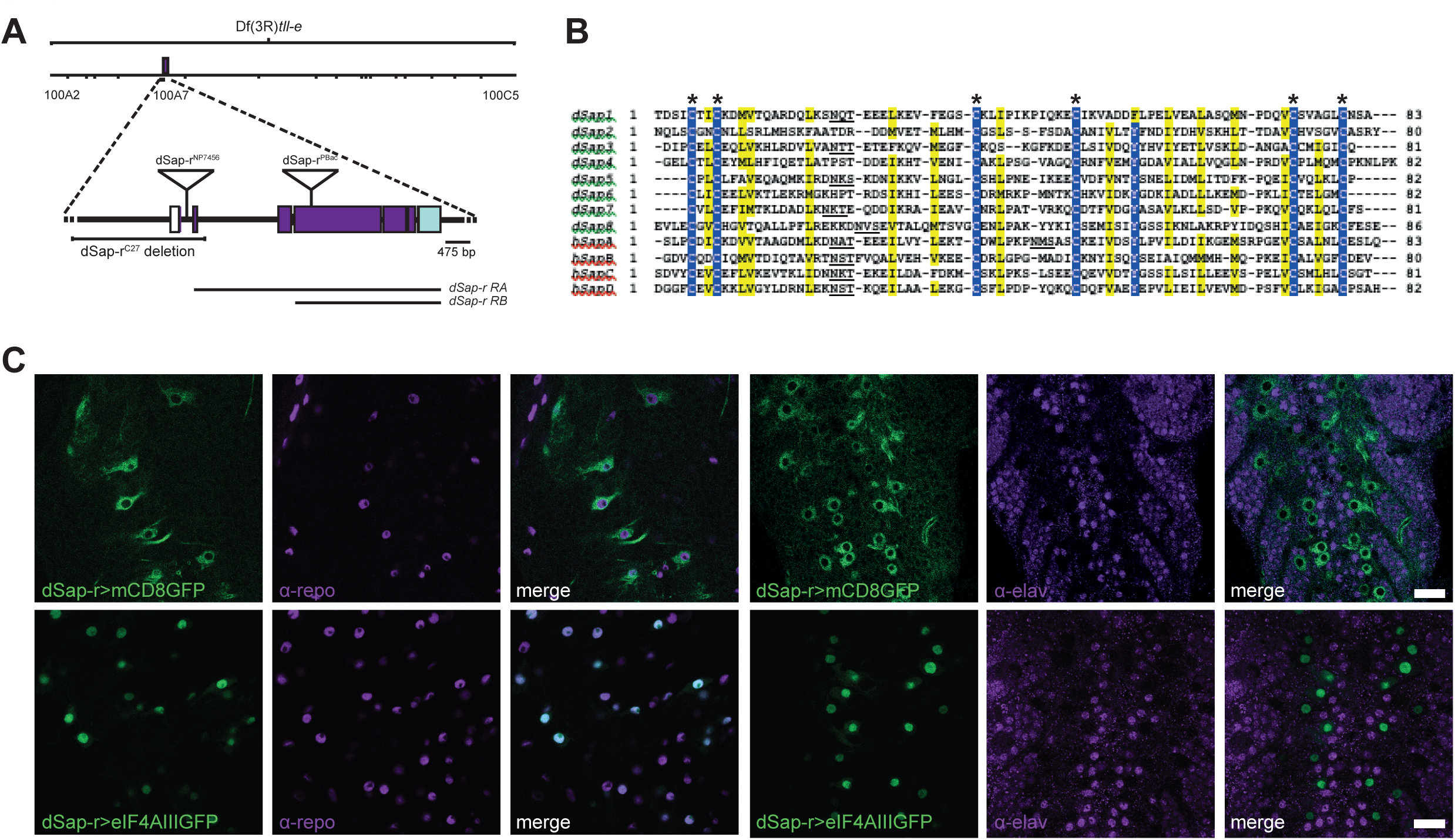
*Drosophila Sap-r* is expressed in glia. A. The *Drosophila prosaposin* homologue *Saposin-related* (*dSap-r*) consists of seven exons and is present on the right arm of the third chromosome (3R) at position 100A7. The *dSap-r* gene contains two potential transcript start sites (ATG), which result in the production of a transcript consisting of the entire coding sequence (*dSap-rRA*, 3451 bp) and an in-frame, truncated version (*dSap-rRB*, 3349 bp) (black lines). The *dSap-r*^*C27*^ allele was generated by *P*-element mobilisation from the *P*{GawB} NP7456 line and is a 2.5kb deletion. The *dSap-r*^*PBac*^ allele consists of a 5.971 bp piggyBac transposon insertion into the largest *dSap-r* exon. The deletion in the deficiency line Df(3R)*tll-e* spans cytogenetic bands 100A2 – 100C5, which includes the *dSap-r* gene. B. A Clustal X alignment of the 8 predicted *Drosophila melanogaster* saposins (dSaps) and the 4 *Homo sapiens* saposins (hSaps). Asterisks, conserved cysteine residues; underlining, glycosylation sites in each hSap and possible homologous sites in the dSaps. Blue highlights identical residues; yellow highlights similar residues (80% threshold setting). C. Confocal images of third instar larvae brains expressing mCD8eGFP or eIF4AIIIGFP under the control of *dSap-r*^*NP7456*^ GAL4 and stained with ±-repo (glial nuclear marker) or ±-elav (pan neuronal marker). Scale bars: 20 µm.

A bl2seq alignment of each human monosaposin (Sap A-D) with the dSap-r protein revealed eight homologous *Drosophila* saposins (dSaps 1-8). Each dSap contained the six conserved cysteine residues critical for saposin function (Fig. 1B). Five of the dSaps also contained a potential glycosylation signal between the second and third cysteines, previously shown in mammals to be necessary for correct saposin folding and function (21).

Saposin proteins are expressed in the mammalian nervous system and their loss usually causes severe neurodegeneration (22-24). We therefore investigated the expression pattern of dSap-r in the nervous system. The *dSap-r*^*NP7456*^ GAL4 enhancer-trap insertion was used to drive a membrane localised GFP reporter (mCD8GFP) in a dSap-r-specific expression pattern. Third instar larvae were dissected and stained with either a glial or neuronal antibody (α-repo or α-elav, respectively). The membrane GFP reporter localised around repo-positive nuclei, suggesting that *dSap-r* is expressed in glial cells (Fig. 1C). The GFP reporter was also shown to faintly surround elav-positive neuronal nuclei, however this is likely to represent expression in glial cells that surround the neuronal cell bodies. To confirm whether *dSap-r* is expressed in neurons, a nuclear GFP reporter (eIF4AIII:GFP) was driven by the *dSap-r*^*NP7456*^ element. No colocalisation was found between the nuclear GFP reporter and neuronal nuclei, suggesting that neurons have no or very little *dSap-r* expression (Fig 1C). The nuclear GFP colocalised with the glial nuclear marker, further confirming *dSap-r* expression in glia.

In mammalian visceral organs, *prosaposin* is expressed ubiquitously at low levels; however moderate to high expression levels are found in the jejunum and tubular epithelial cells in the kidney cortex, epithelial cells of the oesophagus, pancreatic duct and bile duct, and the hepatocytes of the liver (22). PSAP has also been shown to promote spermiogenesis and fertility (25, 26). Using the mCD8GFP reporter in conjunction with the *dSap-r*^*NP7456*^ enhancer-trap element, *dSap-r* was shown to be highly expressed in the reproductive system, the digestive system, Malpighian tubules (*Drosophila* kidney), and the fat bodies (*Drosophila* liver and adipose tissue) of the adult fly (Fig. S1).

The similar expression pattern of dSap-r and mammalian saposins in cells of the nervous, reproductive, digestive, and renal systems, and the liver is suggestive of conserved functions, and therefore supports the use of *Drosophila* to model these disorders.

### *dSap-r* mutation causes a reduced longevity and age-dependent deterioration of locomotion

Patients with saposin deficiency die prematurely, usually within the first decade of life. To test whether *dSap-r* mutants also die prematurely, we first generated deletions of the dSap-r locus via an imprecise P-element mobilisation strategy. The *dSap-r*^*C27*^ allele is a deletion of the first two exons of the *dSap-r* locus (Figure 1A). We also identified a PiggBac transposon insertion, *dSap-r*^*PBac*^, in the 4^th^ exon of the dSap-r locus (Figure 1A). To assess the effect of each *dSap-r* mutation on *dSap-r* transcript levels, RT-PCR was performed on transheterozygous combinations of these two alleles, and combinations of these alleles in trans to a deficiency chromosome uncovering the *dSap-r* locus (Fig. 1A and 2C). The *dSap-r* transcript was almost undetectable in *dSap-r^C27^*/Df mutants and is likely to reflect *dSap-rRB* levels, as the *dSap-rRA* start site is deleted in this mutant. In contrast, *dSap-r* transcript levels in the *dSap-r^PBac^* mutant were indistinguishable from wild type; however, the *dSap-r* primers were designed to anneal upstream of the *dSap-r*^*PBac*^ insertion. Production of saposins from the dSap-r locus prior to the transposon insertion is therefore possible. Using our *dSap-r* mutants, a longevity assay was performed. Each of the three *dSap-r* allelic combinations caused a reduction in longevity compared to wild type flies (Fig. 2A). Median survival for wild type flies was approximately 35 days, whereas 50% of *dSap-r* mutants survived only 6 days (*dSap-r*^*C27*^/Df), 15 days (*dSap-r*^*C27*/*PBac*^) or 18 days (*dSap-r*^*PBac*^/Df). This suggests that the *dSap-r*^*C27*^ allele is more severe than the *dSap-r*^*PBac*^ allele. Taken together with the RT-PCR results, this suggests that the *dSap-r*^*C27*^ allele is likely a strong loss-of-function mutation, whereas the *dSap-r*^*PBac*^ allele may produce dSap-r protein with reduced or aberrant function.

**Fig. 2.**
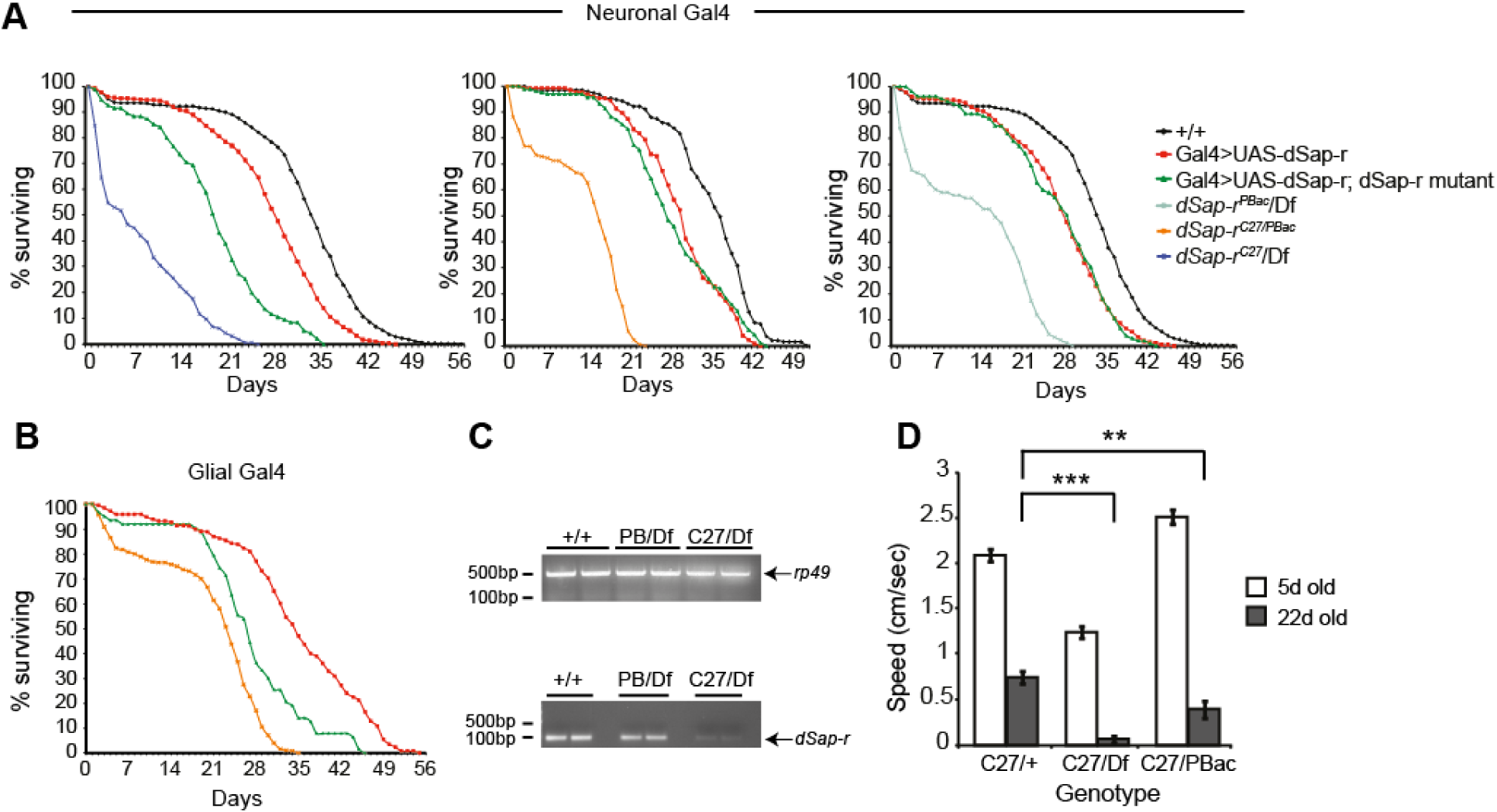
Reduced longevity and physiological deterioration in *dSap-r* mutants. A & B. Longevity of flies with different *dSap-r* allelic combinations was assessed at 29°C. Rescue of *dSap-r* longevity was performed by expressing *dSap-r* cDNA in the neurons (A) or glia (B) of *dSap-r* mutants using the 1407- and repo-GAL4 drivers. Neuronal or glial expression of *dSap-r* in a wild type background was performed as a transgene control. Longevity was plotted as the percentage of Day 0 flies surviving for each genotype. n > 60 for each genotype. C. cDNA from wild type (+/+), *dSap-r*^*PBac*^/Df (PBac/Df) and *dSap-r*^*C27*^/Df (C27/Df) third instar larvae were amplified using *rp49* control and *dSap-r* primers. D. 5-day-old (white bars) and 22-day-old flies (grey bars) were subjected to climbing assays. The climbing speed of *dSap-r*^*C27*^ heterozygotes (C27/+), *dSap-r*^*C27*^/Df (C27/Df) and *dSap-r*^*C27/PBac*^ (C27/PBac) mutants are shown. Error bars: ± sem. * p< 0.05, ** p< 0.005 and *** p≤ 0.001.

To confirm that the *dSap-r* mutations were responsible for the reduced longevity, flies carrying a UAS-*dSap-r* transgene were generated to allow GAL4-driven rescue of *dSap-r* longevity. Ubiquitous expression of *dSap-r* using either Act5c-GAL4 or tubulin-GAL4 resulted in lethality prior to third instar; we therefore used *dSap-r* expression driven by the neuronal 1407- GAL4 (Fig. 2A) or the glial repo-GAL4 (Fig. 2B), either of which did not induce lethality, for rescue experiments. Expression of *dSap-r* in neurons of all *dSap-r* mutant combinations resulted in a substantial rescue of longevity and in most cases longevity was equivalent to the transgene control. Glial *dSap-r* expression was driven in the *dSap-r*^*C27/PBac*^ mutants only, due to greater genetic ease of repo-GAL4 recombination using the *dSap-r*^*PBac*^ allele. Glial *dSap-r* expression also provided a substantial rescue of *dSap-r^C27/PBac^* longevity, however, not to the same degree as the neuronal 1407-GAL4 expression. These results confirm that the reduced longevity in *dSap-r* mutants was due to *dSap-r* mutations.

During the longevity analysis it was observed that the *dSap-r* flies showed an age-dependent decline in locomotion; the 20+ day-old *dSap-r* mutants rarely climbed the vial during spontaneous activity. To quantify this loss of climbing behaviour, 5-day and 22-day-old female flies were tapped to the bottom of a measuring cylinder and their climbing response was captured by video for 45 seconds. Calculation of climbing speed revealed that the 22-day-old *dSap-r*^*C27*^/Df mutants were only able to climb at 5% of their 5-day-old speed compared to maintenance of 36% of 5-day climbing speed for the *dSap-r*^*C27*^/+ controls. The *dSap-r*^*C27/PBac*^ mutants showed an intermediate phenotype maintaining 16% of their 5-day climbing ability at 22-days old. These data confirm an age-dependent deterioration of climbing behaviour in the *dSap-r* mutants (Fig. 2D).

### Age-dependent neurodegeneration in *dSap-r* mutants

The reduced longevity and age-dependent deterioration of locomotion in *dSap-r* mutants is suggestive of progressive neurodegeneration. In *Drosophila*, vacuolarisation of the brain is a hallmark of neurodegeneration (27, 20), which can be quantified from 1 µm tissue sections at the light microscopy level. Horizontal sections were taken from 5-day and 22-day-old fly heads and stained with toluidine blue. Vacuolarisation was specifically observed in regions of sensory function, mainly the antennal lobes, the eye and optic lobes (Fig. 3A). Vacuole number was quantified for these sensory regions (Fig. 3B & C) where we observed that vacuole number in most 5-day-old *dSap-r* mutants was not significantly different from controls, with the exception of limited vacuolarisation observed in the antennal lobes of the *dSap-r*^*C27*^/Df mutants. As the flies aged, the number of vacuoles increased in all *dSap-r* mutants, indicative of progressive neurodegeneration. In the visual system, vacuole number increased by 4 – 18 fold in the *dSap-r* mutants compared to less than a 1.5 fold increase in the controls. In the olfactory system (antennal lobes), the controls showed a modest 1.5 – 3-fold increase in vacuole number. In contrast, the *dSap-r*^*PBac*^/Df and *dSap-r*^*C27/PBac*^ mutants showed a massive 6-fold and 16-fold increase in vacuole number, respectively. Although the *dSap-r*^*C27*^/Df mutants showed the greatest number of vacuoles at 22-days old, the increase was only 3-fold due to the increased severity of vacuolarisation at 5-days old.

**Fig. 3.**
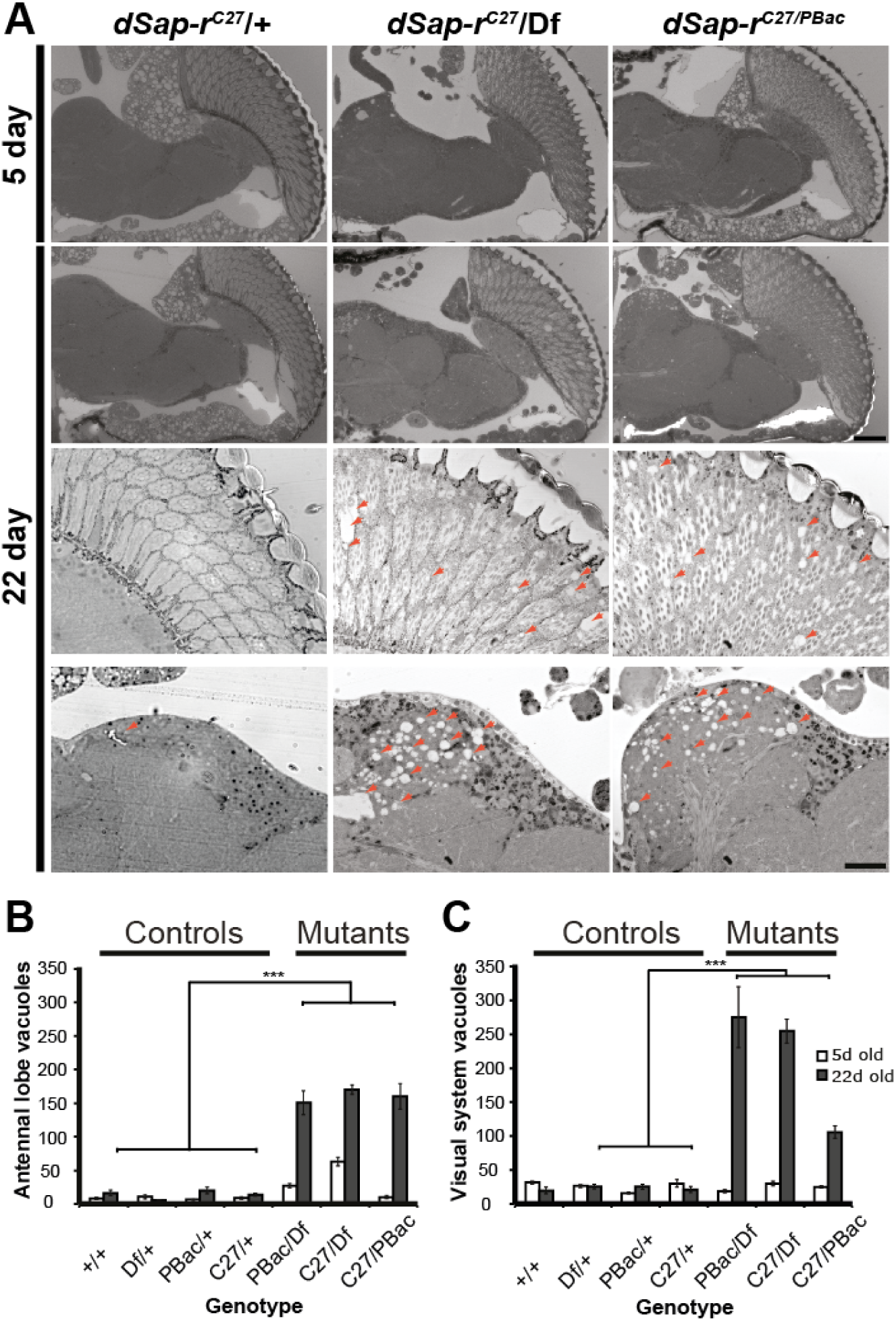
Progressive neurodegeneration in sensory regions of *dSap-r* mutants. A. Representative images of 1 µm sections through 5-day-old and 22-day-old control (dSap-r^C27/^+) and *dSap-r* mutant (*dSap-r*^*C27*^/Df and *dSap-r*^*C27/PBac*^) heads are shown at 20x magnification (top 2 rows) and 63x magnification (bottom 2 rows, eye (upper row) and antennal lobe (lower row)). Scale bars: 100 µm. B & C. Quantification of vacuole number was performed on 3 serial sections per fly (n≥3 flies). Vacuoles were counted for the visual (B; eye, lamina, medulla and lobula complex) and olfactory (C; antennal lobes) systems in both sides of the brain. Quantification is shown for all controls and *dSap-r* mutants tested. Genotypes tested: +/+ (wild type); Df/+, PBac/+ and C27/+ (*dSap-r* heterozygotes); PBac/Df, C27/Df and C27/PBac (*dSap-r* mutants). Error bars: ± sem. *** p≤ 0.001

### *dSap-r* mutants have age-dependent lysosomal storage defects

LSDs are characterised by lysosomal dysfunction leading to the storage of undegraded material in the lysosomes (28). Therefore, to assess the degree of lysosomal dysfunction and storage in *dSap-r* mutants, western blotting was performed using two lysosomal antibodies: anti-Arl-8 and anti-Cathepsin-L.

Arl-8 is an Arf-like GTPase that localises to the lysosomes (29). The Arl-8 antibody was used to probe a western blot containing soluble protein from 5-day and 22-day-old flies to investigate the degree of lysosomal storage in *dSap-r* mutants (Fig. 4A). In 5-day-old flies, Arl-8 levels were similar between wild type and *dSap-r* mutants. However, in 22-day-old *dSap-r* flies, Arl-8 levels were increased compared to wild type and the 5-day-old samples. This suggests an age-dependent accumulation of lysosomal material in *dSap-r* mutants and implies that lysosomes are more abundant and/or swollen in *dSap-r* mutants, which is consistent with the human disease.

**Fig. 4.**
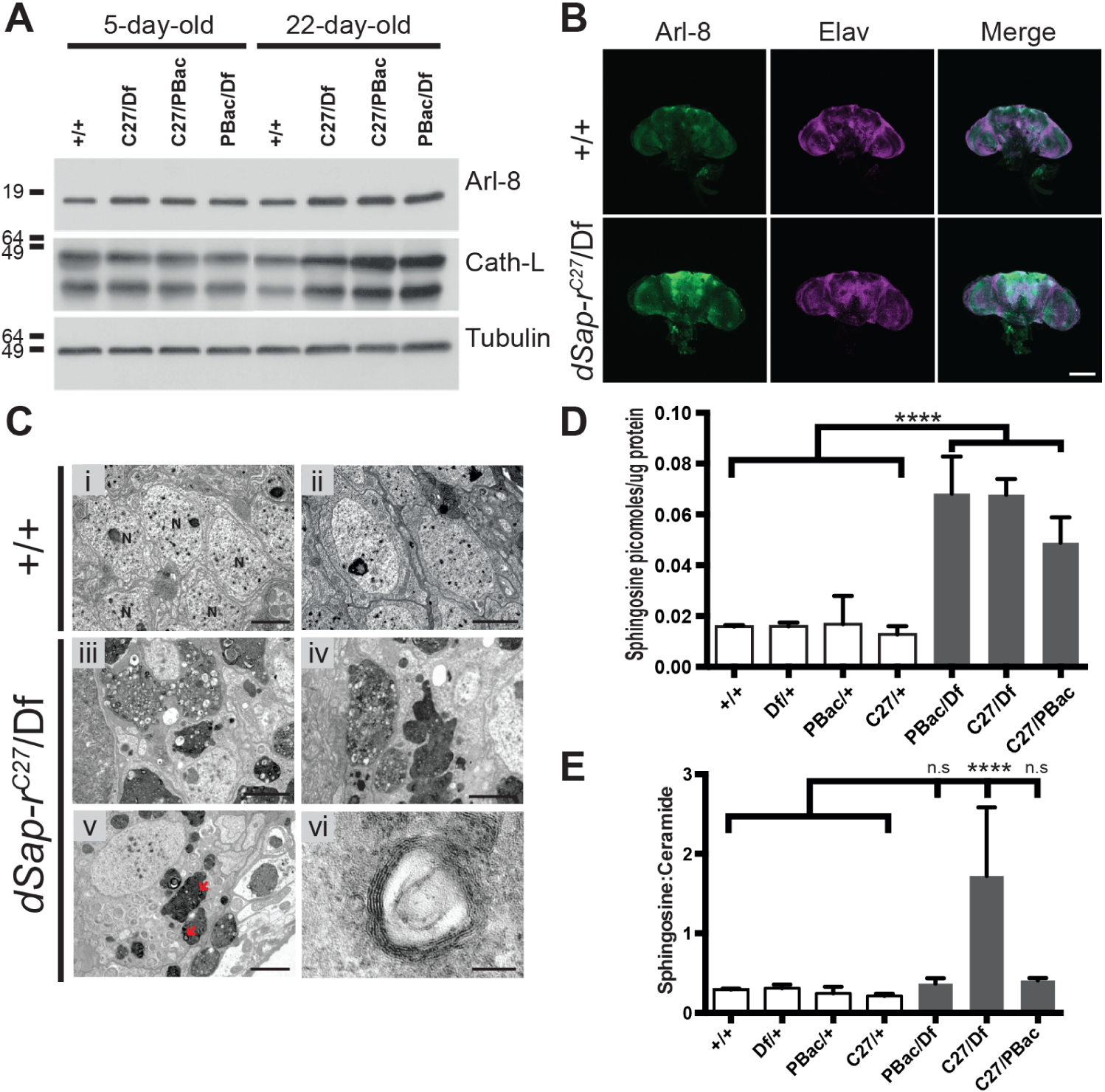
Progressive lysosomal storage in *dSap-r* mutants. A. Western blots showing the abundance of the lysosomal proteins Arl-8 and Cathepsin-L in 5-day-old and 22-day-old wild type (+/+), *dSap-r*^*C27*^/Df (C27/Df), *dSap-r*^*C27*/PBac^ (C27/PBac) and *dSap-r*^*PBac*^/Df (PBac/Df) mutants. Tubulin abundance is show as a loading control. B. Confocal micrographs of 22-day-old wild type (+/+) and *dSap-r*^*C27*^/Df mutant brains showing Arl-8 localisation and abundance. Elav staining shows the similar orientation of both wild type and *dSap-r*^*C27*^/Df mutant brains. Scale bar: 100 μm. C. Transmission electron micrographs of 22-day-old wild type (+/+) and *dSap-r*^*C27*^/Df mutant neuronal cell bodies surrounding the antennal lobes. Multivesicular bodies and multilamellar bodies (arrows) are abundant in *dSap-r*^*C27*^/Df mutant neuronal cell bodies. Scale bars: 2 μm (i - v), 100 nm (vi). D & E. Quantification of sphingosine levels (D) and the sphingosine:ceramide ratio in 5-day-old controls and *dSap-r* mutant brains. Genotypes tested: +/+ (wild type); Df/+, PBac/+ and C27/+ (*dSap-r* heterozygotes); PBac/Df, C27/Df and C27/PBac (*dSap-r* mutants). * p<0.05, *** p<0.001 and **** p<0.0001.

To determine whether this increased abundance of Arl-8 is suggestive of general lysosomal dysfunction, western blot analysis was repeated using an antibody against a lysosomal enzyme (Cathepsin-L). Like all Cathepsins, Cathepsin-L is synthesised as an inactive precursor that is cleaved in the lysosome to form its mature, active form (30). Therefore, if lysosomal function is disturbed in *dSap-r* mutants, Cathepsin-L processing should be less efficient leading to an accumulation of the unprocessed, larger form. Western blot analysis of 5-day and 22-day-old lysates revealed an increase in both the unprocessed and processed forms of Cathepsin-L in all *dSap-r* mutants compared to age-matched controls, but no change in the ratio of unprocessed:processed Cathepsin-L (Fig. 4A). This supports the lysosomal storage phenotype shown by Arl-8 western analysis, but suggests that lysosomal function has not been completely perturbed.

In addition to western blotting, the Arl-8 antibody was used for immunohistochemistry on wild type and *dSap-r*^*C27*^/Df adult brains. An overall increase in fluorescence was observed in the aged *dSap-r* brains, particularly in central brain regions and the antennal lobes (Fig. 4B). Together with our light microscopical analysis of brain sections, this reveals that regions of severe neurodegeneration coincide with the regions of the CNS showing the most lysosomal storage.

A common characteristic of LSDs is the accumulation of membranous storage material known as multilamellar bodies (MLBs) and multivesicular bodies (MVBs). We investigated whether the *dSap-r*^*C27*^/Df mutants carried these hallmark signs. Due to the severe degeneration occurring around the antennal lobes (Fig. 3A), this region of the brain was the main focus for ultrastructural analyses. Electron micrographs of 22-day-old *dSap-r*^*C27*^/Df neurons revealed an abundance of electron dense storage material that has a complex morphology; some regions were populated by electron-lucent droplets, some contained variable numbers of MLBs, and others were densely packed with MVBs (Fig. 4C). The ultrastructural nature of storage material in saposin deficiency patients and mammalian models has been shown to be highly variable, with the presence of MLBs, MVBs and electron-lucent droplets (11, 12, 31, 32). The ultrastructural pathology observed in the *dSap-*r mutants is therefore consistent with the reported disease phenotype, which further supports the use of this model for investigating saposin deficiency.

### Sphingolipids accumulate in *dSap-r* mutant brains

Saposin deficiency leads to the accumulation of a complex array of sphingolipids and sphingolipid intermediates. To determine the nature of sphingolipid perturbation in *dSap-r* mutants, we monitored Hexosyl-Ceramides (HexCer), Ceramide phosphorylethanolamine (CerPE), Ceramide (Cer) and Sphingosine (the breakdown product of Ceramide; 33) in conjunction with a lipidomic analysis of all major membrane phospholipids in the brain (Supplemental Table 1).

Brains from 5-day old flies revealed a consistent increase in sphingolipids (Supplemental Table 1), particularly sphingosine, in all *dSap-r* mutants examined (Fig. 4D). Notably, comparison of the sphingosine:ceramide ratios in controls and mutants revealed a striking imbalance between these sphingolipid intermediates in the *dSap-r*^*C27/Df*^ mutants (Fig. 4E). We do not observe a significant change in HexCer across the mutant combinations (Supplemental Table 1).

Changes in particular phospholipid classes were observed. However, unlike the sphingolipids, there is no specific and consistent trend for changes in phospholipid classes across the different *dSap-r* mutant combinations tested (Supplemental Table 1).

### *dSap-r*^*C27*^/Df CNS soma are enlarged with storage

During ultrastructural analyses of the CNS we observed that *dSap-r^C27^*/Df neuronal soma were consistently and grossly enlarged compared to wild type. To quantify this difference, the soma boundaries of each cell were marked using ImageJ and the soma area measured (Fig. 5A). Measuring the average soma area revealed an almost 2.5-fold increase in *dSap-r*^*C27*^/Df mutants compared to wild type (Fig. 5B). When normalised to nucleus area, the *dSap-r*^*C27*^/Df soma were almost 2-fold larger suggesting that nuclear area is also increased in *dSap-r*^*C27*^/Df neurons (Fig. 5C). We also observed cell enlargement and increased storage in glial cells of the *dSap-r*^*C27*^/Df adult CNS (Supplemental Fig. 2).

**Fig. 5.**
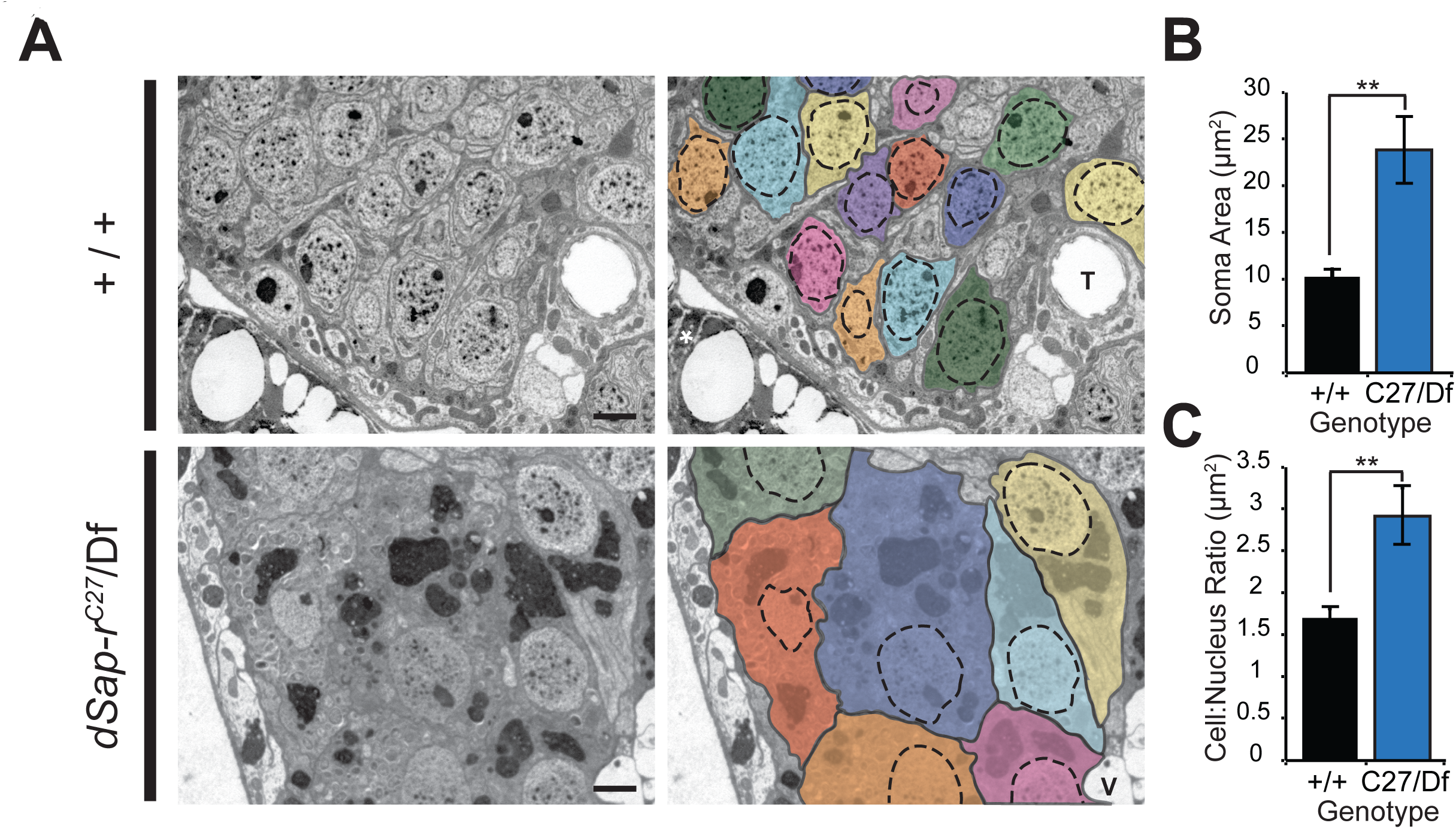
*dSap-r* mutant brains have grossly enlarged soma size. A. Transmission electron micrographs showing neuronal cell bodies adjacent to the antennal lobes of 22-day-old wild type (+/+) and *dSap-r*^*C27*^/Df mutants. Each soma has been demarcated in grey and rendered a different pseudocolour. The nuclei are demarcated by dashed lines. T, trachea; V, vacuole. Scale bar: 2 μm. B & C. Quantification of 22-day-old wild type (+/+) and *dSap-r*^*C27*^/Df (C27/Df) soma area (B) and cell:nucleus ratio (C). ** p<0.005.

### Loss of *dSap-r* function causes a deterioration of visual function

We have shown that sensory regions of the brain are particularly vulnerable to *dSap-r* mutation. To assess the effect of *dSap-r* mutation on the function of sensory neurons, we investigated the integrity of the visual system in *dSap-r* flies. Electron micrographs revealed an abundance of stored material in photoreceptor neurons of the fly retina (Fig. 6A), similar in morphology to that found in the antennal lobe neurons (Fig. 4C). High magnification images showed a variable phenotype in the rhabdomeres, the light responsive component of the eye. Most rhabdomeres showed massive accumulation of electron-dense material yet some were found to be structurally intact (Fig. 6A).

**Fig. 6.**
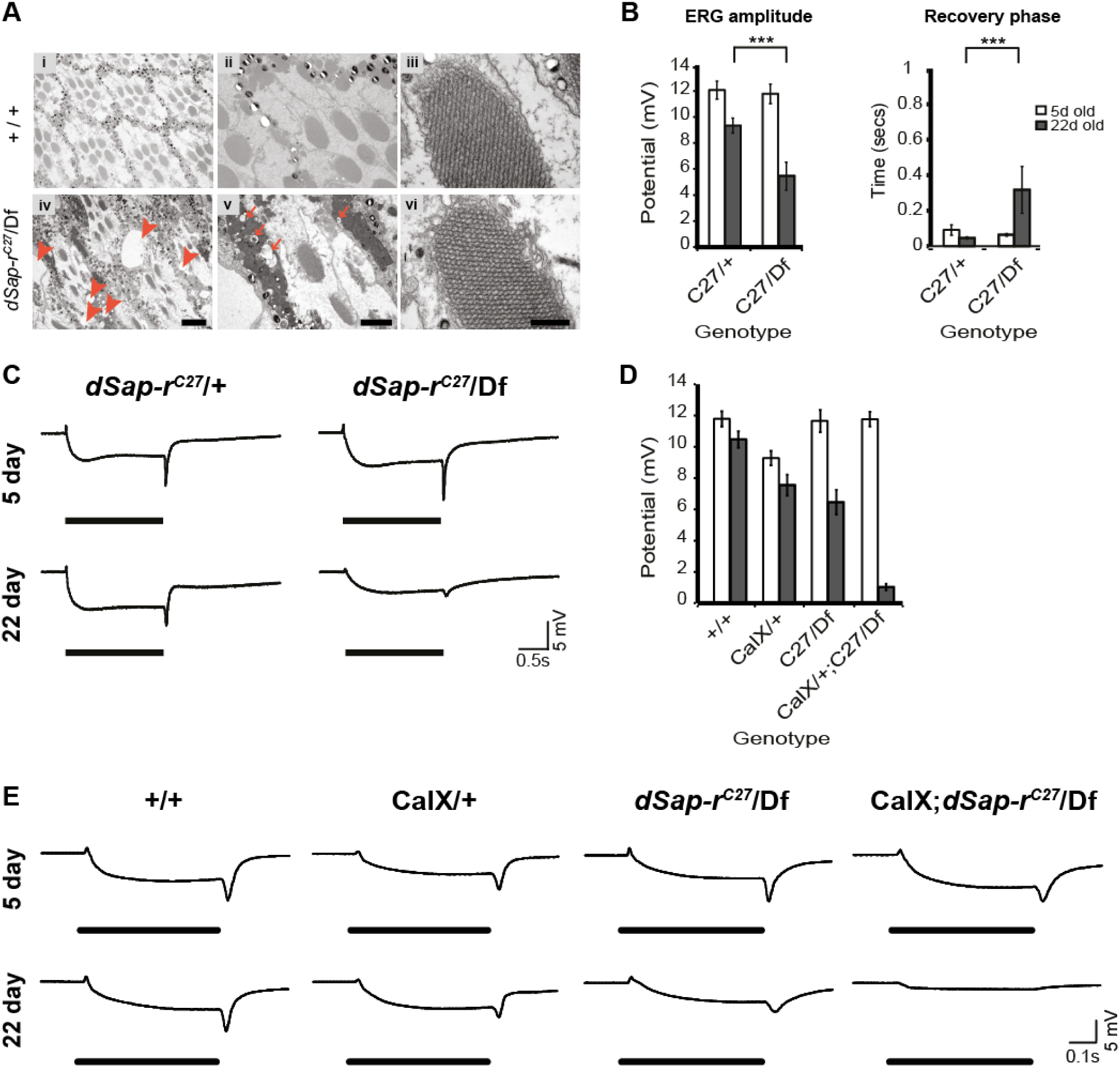
Calcium homeostasis defects are associated with progressive deterioration of visual function in *dSap-r* mutants. A. Transmission electron micrographs showing the integrity of the ommatidia in 22-day-old wild type (+/+) and *dSap-r*^*C27*^/Df mutants. Arrowheads mark vacuoles and arrows mark electron lucent material within regions of electron-dense storage. Scale bars: 5 μm (i & iv), 2 μm (ii & v) and 500 nm (iii & vi). B. Quantification of electroretinogram (ERG) amplitude and recovery rate after a blue light pulse. The recovery rate is the time taken for the potential to reach half way between the off-transient and base-line potentials after termination of the light pulse. C. Representative ERG traces for 5-day-old and 22-day-old control (*dSap-r*^*C27*^/+) and *dSap-r*^*C27*^/Df mutant females following a blue light pulse. D. Quantification of ERG amplitude after a blue light pulse in 5-day-old and 22-day-old wild type overexpressing the calcium exchanger CalX in the eye (CalX/+), *dSap-r*^*C27*^/Df mutants (C27/Df) and *dSap-r*^*C27*^/Df mutants overexpressing CalX in the eye (CalX/+;C27/Df). E. Representative ERG traces for 5-day-old and 22-day-old wild type overexpressing CalX (CalX/+), *dSap-r*^*C27*^/Df mutants and *dSap-r*^*C27*^/Df mutants expressing CalX (CalX; *dSap-r*^*C27*^/Df). N ≥ 7 flies per condition. *** p≤0.001

To determine whether the photoreceptor neurons and their underlying optic lobe synapses were functionally intact, we measured the ability of the fly eye to respond to light using the electroretinogram (ERG). The ERG is a classic method for investigating photoreceptor and lamina neuron function by measuring the summed potential difference on the surface of the fly eye in response to light pulses (34, 35). Different components of the ERG trace represent the net influence of different parts of the visual system: the initial decline in potential reflects the depolarisation of the photoreceptor neurons; the on- and off-transients are a result of synaptic transmission with the L1 and L2 lamina neurons; and the recovery phase denotes the feedback from the optic lobe to the photoreceptor neurons required for efficient repolarisation of the photoreceptors (34–37). Therefore, the integrity of the different components of visual transduction can be assessed using this simple approach.

ERG recordings in 5-day-old *dSap-r*^*C27*^/Df mutants were not significantly different from controls in all components measured (Fig. 6B & C). However, 22-day-old *dSap-r*^*C27*^/Df mutants showed a severe deterioration of all components of the ERG compared to controls. The *dSap-r*^*C27*^/*Df* ERG potential deteriorated to 46% of its 5-day value, whereas the *dSap-r*^*C27*^/+ control ERG potential only decreased to 77% of its 5-day value. This severe deterioration of the ERG amplitude in *dSap-r*^*C27*^/*Df* mutants was also matched by a 5-fold increase in recovery time after termination of the light pulse. This suggests that neuronal function in both the retina and underlying optic lobe is severely compromised in *dSap-r* mutants.

### Calcium homeostasis is defective in *dSap-r* mutants

Calcium homeostasis defects have been highlighted as one aspect of cellular pathology implicated in the sphinoglipidoses (38-40). For example, in NPC1 cells lysosomal calcium levels have been shown to be abnormally low, and by increasing cytosolic calcium levels NPC pathology in both NPC1 cells and mouse models can be substantially rescued (40). Calcium homeostasis defects have also been implicated in other sphingolipidoses (Gaucher’s, Sandhoff and GM_1_-gangliosidosis); however, an increase in cytosolic calcium levels was pathological in these cases (41-44).

To investigate if the *dSap-r* sphingolipidosis model showed a calcium homeostasis defects, we measured ERGs in *dSap-r* mutants overexpressing the plasma membrane calcium exchanger (CalX) in the fly eye using the *NINAE* (Rhodopsin 1) promoter. CalX transports calcium ions out of the neuron in exchange for sodium ions (45). Therefore, if *dSap-r* mutants have a reduced cytosolic calcium level, CalX expression should exacerbate the degeneration and function of the photoreceptors; however, the ERG would be substantially rescued if *dSap-r* mutation causes an increased cytosolic calcium level.

Overexpression of CalX in an otherwise wild type fly resulted in a modest degeneration of the ERG to 81% of its 5-day level. When CalX was expressed in a *dSap-r*^*C27*^/Df background, there was no effect on the 5-day-old *dSap-r*^*C27*^/Df ERG. However, as the flies aged, CalX expression induced a decrease in *dSap-r*^*C27*^/Df vision to only 9% of its 5-day-old level; this is in contrast to the deterioration of the *dSap-r*^*C27*^/Df ERG to 64% of its 5-day-old level, and therefore suggests that the *dSap-r* mutation may cause an abnormally reduced cytosolic calcium level.

## Discussion

To further our understanding of saposin deficiency disease, we generated a *Drosophila* model. The *Drosophila* prosaposin homologue *dSap-r* was revealed to contain eight saposin-like domains, which all contained the six-cysteine primary sequence common to all saposins (this investigation, 46). We showed that most of the dSap peptides contain a predicted glycosylation site, a feature critical for saposin folding and function (21). The sequence similarities between the mammalian and *Drosophila* saposins suggested that we had identified the correct ortholog.

To further characterise dSap-r, we investigated its expression pattern using GFP reporters. Both membrane and nuclear GFP reporters confirmed the expression of *dSap-r* in glia, as reported by (47). Our findings also suggested that, unlike in mammals, *dSap-r* is either not expressed in neurons or is expressed at low/undetectable levels.

Although *dSap-r* expression has been shown to occur in glia, this does not discount the presence of dSap-r protein in neurons. Mammalian prosaposin is abundant in secretory fluids, including cerebrospinal fluid, seminal fluid, milk, bile and pancreatic fluid (48-51); in fact prosaposin is one of the main secretory products of Sertoli cells in the male reproductive system (52). Therefore, we propose that *Drosophila* glia may provide dSap-r to neurons by glial secretion and neuronal uptake. This notion is supported by our findings that *dSap-r* longevity can be substantially rescued by expressing *dSap-r* directly in neurons, which suggests a neuronal requirement for dSap-r. This is further supported by our TEM analyses showing more severe lysosomal storage and consequential increase in cell size of the neurons compared to the glia. This non-autonomous function of dSap-r has important implications for potential therapeutic strategies as it suggests that a source of prosaposin secreting cells could provide a successful intervention for these conditions.

This investigation also revealed high *dSap-r* expression in the male and female reproductive organs, the digestive system, *Drosophila* renal system (Malpighian tubules), and the *Drosophila* liver equivalent (the fat bodies). Mammalian prosaposin has a role in spermiogenesis and improving fertility (25, 26, 53, 54), and is also relatively abundant in subsets of cells of the small intestine, kidney and liver (22). Therefore, in addition to having a conserved primary sequence, dSap-r also has a conserved tissue expression suggesting a conserved function.

After confirming the identity of the *Drosophila* prosaposin orthologue and its expression pattern, we generated a *dSap-r* loss-of-function model and assessed its pathology. Ultrastructural analysis revealed extensive storage in the *dSap-r* nervous system (MLBs, MVBs and lipid droplets), characteristic of LSD pathology. Lysosomal storage was also confirmed by western blotting, which showed progressive accumulation of two lysosomal markers (Arl8 and Cathepsin-L) in the *dSap-r* mutants, and mass spectrometry revealed early accumulation and imbalance of sphingolipid intermediates in *dSap-r* mutants. The *dSap-r*^*C27/Df*^ mutants were the most severe mutants in all assays tested. They also showed the most severe imbalance of sphingosine and ceramide, whilst having similar levels of sphingosine and lower levels of ceramide than the other mutant allelic combinations. This therefore supports previous findings that the ratio of sphingolipids is more important than the total levels of individual sphingolipids when considering the effect on pathology (54-58).

In prosaposin and individual saposin deficiency mouse models, the stored material was described as very heterogeneous with electron-dense and electron-lucent inclusions, MLBs and MVBs (32, 54, 56, 58-61). This phenotype is strikingly similar to the ultrastructural pathology of *dSap-r* mutants.

Like other *Drosophila* LSD models, *dSap-r* mutants die prematurely, with 50% death within 18 days; this equates to a 41-49% reduction in longevity, which is comparable to most *Drosophila* LSD models (40-55% reduction) (19, 20, 62-64). The *dSap-r* mutants also showed an age-dependent deterioration of spontaneous and evoked locomotion, which correlated with the progressive nature of vacuolarisation in the sensory regions of the *dSap-r* nervous system. This may suggest that locomotion deterioration is primarily a result of failed perception of sensory cues rather than motor neuron or muscle degeneration. This notion is supported by a less severe deterioration in *dSap-r* jump performance using an assay that bypasses sensory neuron input (data not shown). A similar degeneration in sensory function is observed in a *Drosophila* model of NPC1 (20) and more recently recognised in NPC1 patients (73). These results suggest a particular sensitivity of sensory neurons to disruptions in sphingolipid metabolism. This is supported by the severe sensory degeneration observed in the sphingolipid synthesis disorder Hereditary Sensory and Autonomic Neuropathy Type 1 (HSAN1) (65). HSAN1 is caused by mutations in the first enzyme of the sphingolipid synthesis pathway: serine-palmitoyl transferase (SPT). Although HSAN1 is a sphingolipid synthesis disorder and saposin deficiency is a disorder of sphingolipid degradation, the sensory degeneration in both of these disorders reflects a common cellular pathology: a defect in sphingolipid homeostasis.

Therefore, sensory degeneration may result from an absence of sphingolipids, an abnormal accumulation of sphingolipids, an imbalance of sphingolipids, or a combination of all three. Although retinal degeneration and storage has been reported in other *Drosophila* models of LSDs (66 27 20 67), the pathological mechanism remains unclear. In several mammalian sphingolipidoses models, recent reports have revealed calcium homeostasis defects. Therefore, a calcium exchanger was overexpressed in the fly eye of *dSap-r* mutants to determine whether altering cytosolic calcium levels had any effect on *dSap-r* retinal function. Aging of *dSap-r*^*C27/Df*^ mutants caused a 36% reduction in retinal function; a similar but less severe (19%) deterioration was also seen after expression of CalX in aged wild type flies. Overexpression of CalX in a *dSap-r* mutant background caused a severe (92%) deterioration of retinal function, leaving the 22-day-old flies almost completely unresponsive to light. This significant genetic interaction suggests that calcium regulation is compromised in *dSap-r* mutant photoreceptors, leading to neurodegeneration and perturbed sensory function.

The *Drosophila* NPC1 model revealed degeneration of the visual system and concomitant deterioration of the ERG that resembles that of the *dSap-r* model (20). Although classically characterised as a cholesterol-storage disorder, NPC pathology has more recently been linked to sphingolipid storage (40), specifically a sphingosine storage leading to a deficit in lysosomal calcium regulation (40); therefore, calcium homeostasis may also be dysfunctional in the NPC1 *Drosophila* model. In support of this, in cell culture, NPC1 cells contained approximately 65% less lysosomal calcium compared to control cells. It was shown that the sphingolipid intermediate sphingosine was responsible for this calcium homeostasis defect (40), directly linking calcium homeostasis defects to the accumulation of sphingolipids in the sphingolipidoses.

Our sphingolipid analyses showed that sphingosine levels were significantly increased in all *dSap-r* mutant brains. Like in NPC1 cells, this may cause a decrease in lysosomal calcium in *dSap-r* mutants, which was further exacerbated by overexpression of CalX. Further evidence of a link between sphingolipid metabolism and calcium homeostasis was also provided by (68), who revealed that retinal degeneration caused by imbalances in sphingolipid metabolites was suppressed by mutations in the *Drosophila CalX*. We have also shown an imbalance in the ratio between two sphingolipid intermediates in *dSap-r* brains, providing further support of calcium homeostasis defects being a likely cause of *dSap-r* pathology.

In conclusion, we suggest that in *dSap-r* mutants an early accumulation of sphingosine, or the imbalance of the sphingosine:ceramide ratio, leads to a decrease in lysosomal calcium. This calcium deficiency causes degeneration of sensory neurons in the *dSap-r* mutants leading to physiological deterioration. Therapeutic interventions to increase cytosolic calcium levels in saposin-deficient patients may therefore provide a successful route for ameliorating these disorders.

## Funding

This work was funded by a Quota studentship from the BBSRC [to S.J.H] and a Medical Research Council grant [G0400580 to S.T.S]. Work in the D.S laboratory was supported by a Senior Fellowship of the Wellcome Trust/DBT India Alliance, and the NCBS-Merck & Co International Investigator Award [D.S].

## Acknowledgements

For fly stocks and reagents we would like to thank Andreas Prokop, Debbie Smith, Dani Ungar, Roger Hardie, The Bloomington and Kyoto *Drosophila* stock centres and the Developmental Studies Hybridoma Bank, Iowa. The anti-repo, anti-elav and anti-tubulin monoclonal antibodies, developed by Corey Goodman, Gerald M. Rubin and Michael Klymkowsky, were obtained from the DSHB, created by the NICHD of the NIH and maintained at the University of Iowa, Department of Biology, Iowa, IA 52242. For technical advice we are indebted to Meg Stark, Karen Hodgkinson, Graeme Park, Gareth Evans and Paul Pryor.

## Conflict of interest statement

None declared.

## Abbreviations

PSAP: Prosaposin
ERG: electroretinograms
LSD: lysosomal storage disease
NPC: Niemann Pick type C
GFP: green fluorescent protein
RT-PCR: reverse transcription PCR
TEM: transmission electron microscopy
MVB: multivesicular body
MLB: multilamellar body
HSAN1: Hereditary sensory and autonomic neuropathy type 1

## Figure Legends

**Supplemental Fig. 1.**
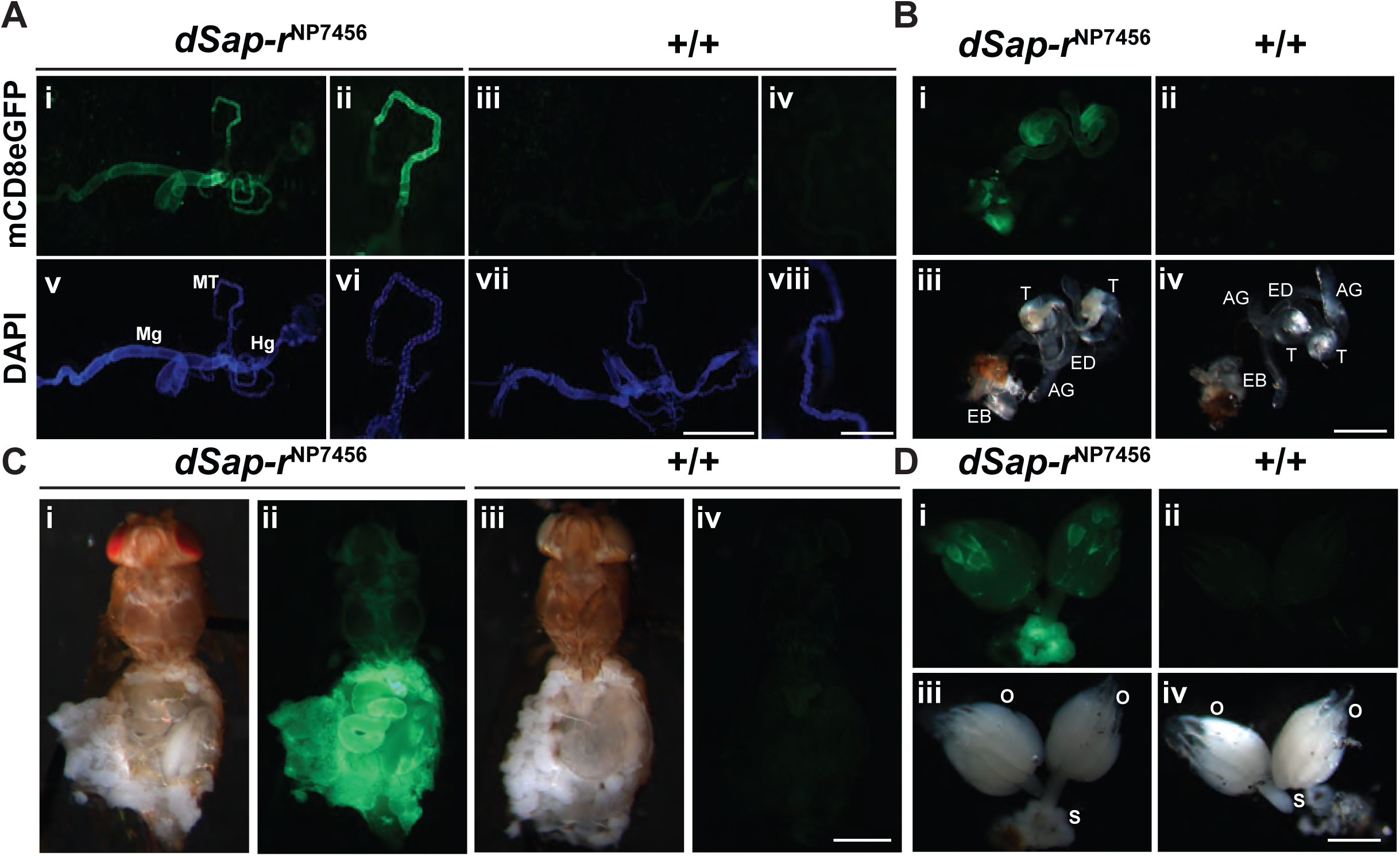
*dSap-r* is expressed in visceral organs of *Drosophila.* Digestive systems (A), male (B) and female (D) reproductive systems and fat bodies (C) are shown from adult controls and flies expressing mCD8eGFP under the control of *dSap-r*^*NP7456*^ GAL4. Organs are stained with the nuclear marker DAPI in (A). MT, Malpighian tubule; Mg, midgut; Hg, hindgut; T, testes; EB, ejaculatory bulb; AG, accessory gland; ED, ejaculatory duct; O, ovary; S, spermatheca. Scale bars: (A) 1000 µm (vii) and 250 µm (viii), (B-D) 500 µm.

**Supplemental Fig. 2.**
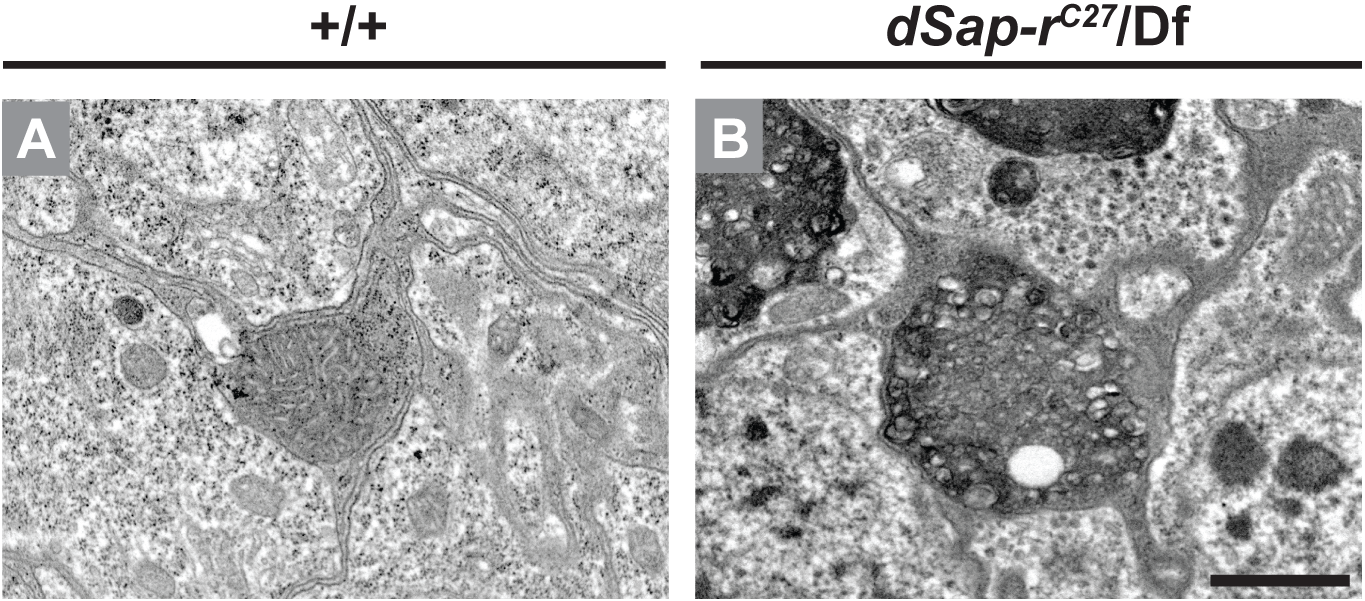
Cell enlargement and increased storage in *dSap-r*^*C27*^/Df glia. Transmission electron micrographs of glia surrounding the antennal lobe of 22-day old wild type (+/+; A) and *dSap-r*^*C27*^/Df (B) brains. Electron-dense and electron-lucent vesicular storage is shown in *dSap-r*^*C27*^/Df glia (B). Scale bar: 1 µm. n = 3

**Supplemental Table 1:**
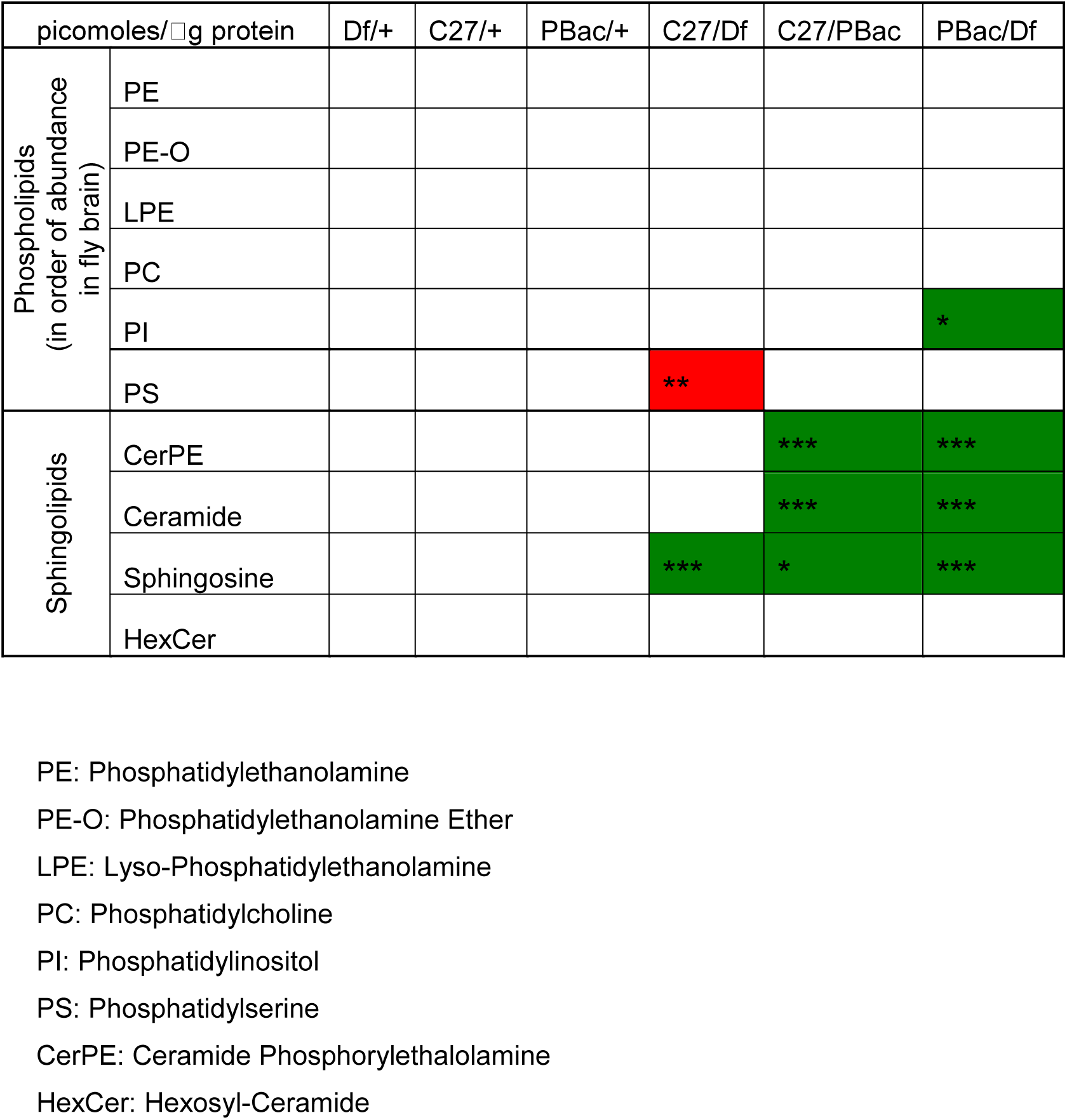
Summary of lipidome-wide changes in *dSap-r* mutant brains. Table depicts the significant changes in levels of major sphingolipids and compared to +/+ (Canton-S; Control). A minimum of 4 replicates were used for the analyses. Statistically significant changes are indicated by green (increase) and red (decrease). * p<0.01, **p<0.001, ***p<0.0001 as determined by ANOVA followed by post-hoc Bonferonni test.

| picomoles/□g protein |  | Df/+ | C27/+ | PBac/+ | C27/Df | C27/PBac | PBac/Df |
| --- | --- | --- | --- | --- | --- | --- | --- |
| Phospholipids<br>(in order of abundance<br>in fly brain) | PE |  |  |  |  |  |  |
|  | PE-O |  |  |  |  |  |  |
|  | LPE |  |  |  |  |  |  |
|  | PC |  |  |  |  |  |  |
|  | PI |  |  |  |  |  | * |
|  | PS |  |  |  | ** |  |  |
| Sphingolipids | CerPE |  |  |  |  | *** | *** |
|  | Ceramide |  |  |  |  | *** | *** |
|  | Sphingosine |  |  |  | *** | * | *** |
|  | HexCer |  |  |  |  |  |  |

PE: Phosphatidylethanolamine
PE-O: Phosphatidylethanolamine Ether
LPE: Lyso-Phosphatidylethanolamine
PC: Phosphatidylcholine
PI: Phosphatidylinositol
PS: Phosphatidylserine
CerPE: Ceramide Phosphorylethanolamine
HexCer: Hexosyl-Ceramide

**Supplemental Table 2:**
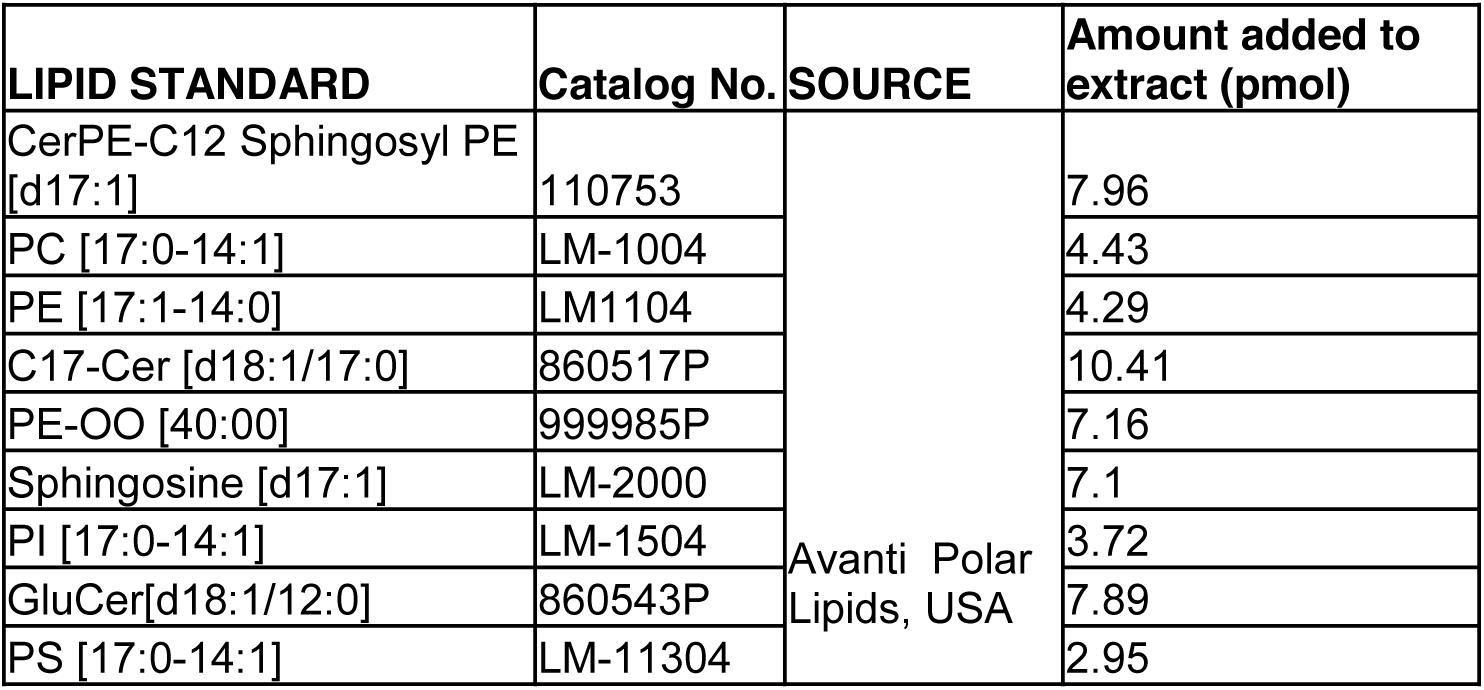
Internal Standard (IS) mix used for lipidomics

## References

1. Furst, W., Machleidt, W. and Sandhoff, K. (1988). The precursor of sulfatide activator protein is processed to three different proteins. Biol. Chem. Hoppe Seyler 369, 317–328.

2. O'Brien, J.S., Kretz, K.A., Dewji, N., Wenger, D.A., Esch, F. and Fluharty, A.L. (1988). Coding of two sphingolipid activator proteins (SAP-1 and SAP-2) by same genetic locus. Science 241, 1098–1101.

3. Nakano, T., Sandhoff, K., Stumper, J., Christomanou, H. and Suzuki, K. (1989). Structure of full-length cDNA coding for sulfatide activator, a Co-beta-glucosidase and two other homologous proteins: two alternate forms of the sulfatide activator. J. Biochem. 105, 152–154.

4. Hiraiwa, M., Martin, B.M., Kishimoto, Y., Conner, G.E., Tsuji, S. and O'Brien, J.S. (1997). Lysosomal proteolysis of prosaposin, the precursor of saposins (sphingolipid activator proteins): its mechanism and inhibition by ganglioside. Arch. Biochem. Biophys. 341, 17–24.

5. Berent, S.L. and Radin, N.S. (1981). Mechanism of activation of glucocerebrosidase by co-beta-glucosidase (glucosidase activator protein). Biochim. Biophys. Acta. 664, 572–582.

6. Vogel, A., Furst, W., Abo-Hashish, M.A., Lee-Vaupel, M., Conzelmann, E. and Sandhoff, K. (1987). Identity of the activator proteins for the enzymatic hydrolysis of sulfatide, ganglioside GM1, and globotriaosylceramide. Arch. Biochem. Biophys. 259, 627–638.

7. Morimoto, S., Martin, B.M., Yamamoto, Y., Kretz, K.A., O'Brien, J.S. and Kishimoto, Y. (1989). Saposin A: second cerebrosidase activator protein. Proc. Natl. Acad. Sci. U S A. 86, 3389–3393.

8. Azuma, N., O'Brien, J.S., Moser, H.W. and Kishimoto, Y. (1994). Stimulation of acid ceramidase activity by saposin D. Arch. Biochem. Biophys. 311, 354–357.

9. Yamada, M., Inui, K., Hamada, D., Nakahira, K., Yanagihara, K., Sakai, N., Nishigaki, T., Ozono, K. and Yanagihara, I. (2004). Analysis of recombinant human saposin A expressed by Pichia pastoris. Biochem. Biophys. Res. Commun. 318, 588–593.

10. Elleder, M., Jirasek, A., Smid, F., Ledvinova, J., Besley, G. T. and Stopekova, M. (1984). Niemann-Pick disease type C with enhanced glycolipid storage. Report on further case of so-called lactosylceramidosis. Virchows Arch. A. Pathol. Anat. Histopathol. 402, 307–317.

11. Harzer, K., Paton, B.C., Poulos, A., Kustermann-Kuhn, B., Roggendorf, W., Grisar, T. and Popp, M. (1989). Sphingolipid activator protein deficiency in a 16-week-old atypical Gaucher disease patient and his fetal sibling: biochemical signs of combined sphingolipidoses. Eur. J. Pediatr. 149, 31–39.

12. Hulkova, H., Cervenkova, M., Ledvinova, J., Tochackova, M., Hrebicek, M., Poupetova, H., Befekadu, A., Berna, L., Paton, B.C., Harzer, K. et al. (2001). A novel mutation in the coding region of the prosaposin gene leads to a complete deficiency of prosaposin and saposins, and is associated with a complex sphingolipidosis dominated by lactosylceramide accumulation. Hum. Mol. Genet. 10, 927–940.

13. Spiegel, R., Bach, G., Sury, V., Mengistu, G., Meidan, B., Shalev, S., Shneor, Y., Mandel, H. and Zeigler, M. (2005). A mutation in the saposin A coding region of the prosaposin gene in an infant presenting as Krabbe disease: first report of saposin A deficiency in humans. Mol. Genet. Metab. 84, 160–166.

14. Tylki-Szymanska, A., Czartoryska, B., Vanier, M.T., Poorthuis, B.J., Groener, J.A., Lugowska, A., Millat, G., Vaccaro, A.M. and Jurkiewicz, E. (2007). Non-neuronopathic Gaucher disease due to saposin C deficiency. Clin. Genet. 72, 538–542.

15. Christomanou, H., Aignesberger, A. and Linke, R.P. (1986). Immunochemical characterization of two activator proteins stimulating enzymic sphingomyelin degradation in vitro. Absence of one of them in a human Gaucher disease variant. Biol. Chem. Hoppe Seyler. 367, 879–890.

16. O'Brien, J.S. and Kishimoto, Y. (1991). Saposin proteins: structure, function, and role in human lysosomal storage disorders. Faseb. J. 5, 301–308.

17. Huang, X., Suyama, K., Buchanan, J., Zhu, A.J. and Scott, M.P. (2005). A Drosophila model of the Niemann-Pick type C lysosome storage disease: dnpc1a is required for molting and sterol homeostasis. Development 132, 5115–5124.

18. Fluegel, M.L., Parker, T.J. and Pallanck, L.J. (2006). Mutations of a Drosophila NPC1 gene confer sterol and ecdysone metabolic defects. Genetics 172(1), 185–196.

19. Huang, X., Warren, J.T., Buchanan, J., Gilbert, L.I. and Scott, M.P. (2007). Drosophila Niemann-Pick Type C-2 genes control sterol homeostasis and steroid biosynthesis: a model of human neurodegenerative disease. Development 134, 3733–3742.

20. Phillips, S.E., Woodruff, E.A., 3rd, Liang, P., Patten, M. and Broadie, K. (2008). Neuronal loss of Drosophila NPC1a causes cholesterol aggregation and age-progressive neurodegeneration. J. Neurosci. 28, 6569–6582.

21. Hiraiwa, M., Soeda, S., Martin, B.M., Fluharty, A.L., Hirabayashi, Y., O'Brien, J.S. and Kishimoto, Y. (1993). The effect of carbohydrate removal on stability and activity of saposin B. Arch. Biochem. Biophys. 303, 326–331.

22. Sun, Y., Witte, D.P. and Grabowski, G.A. (1994). Developmental and tissue-specific expression of prosaposin mRNA in murine tissues. Am. J. Pathol. 145, 1390–1398.

23. Van Den Berghe, L., Sainton, K., Gogat, K., Marchant, D., Dufour, E., Bonnel, S., Gadin, S., Menasche, M. and Abitbol, M. (2004). Prosaposin gene expression in normal and dystrophic RCS rat retina. Invest. Ophthalmol. Vis. Sci. 45, 1297–1305.

24. Yoneshige, A., Suzuki, K., Kojima, N. and Matsuda, J. (2009). Regional expression of prosaposin in the wild-type and saposin D-deficient mouse brain detected by an anti-mouse prosaposin-specific antibody. Proc. Jpn. Acad. Ser. B. Phys. Biol. Sci. 85, 422–434.

25. Hammerstedt, R.H. (1997). A method and use of polypeptide in sperm-egg binding to enhance or decrease fertility. International Patent Publication Number W)/97/25620 Geneva: World International Property Organization, 1–42

26. Amann, R.P., Hammerstedt, R.H. and Shabanowitz, R.B. (1999). Exposure of human, boar, or bull sperm to a synthetic peptide increases binding to an egg-membrane substrate. J. Androl. 20, 34–41.

27. Dermaut, B., Norga, K.K., Kania, A., Verstreken, P., Pan, H., Zhou, Y., Callaerts, P. and Bellen, H.J. (2005). Aberrant lysosomal carbohydrate storage accompanies endocytic defects and neurodegeneration in Drosophila benchwarmer. J. Cell Biol. 170, 127–139.

28. Jardim, L.B., Villanueva, M.M., de Souza, C.F. and Netto, C.B. (2010). Clinical aspects of neuropathic lysosomal storage disorders. J. Inherit. Metab. Dis. 33, 315–329.

29. Hofmann, I. and Munro, S. (2006). An N-terminally acetylated Arf-like GTPase is localised to lysosomes and affects their motility. J. Cell Sci. 119, 1494–1503.

30. Erikson, A.H. (1989). Biosynthesis of lysosomal endopeptidases. J. Cell Biochem. 40(1), 31–41.

31. Fujita, N., Suzuki, K., Vanier, M.T., Popko, B., Maeda, N., Klein, A., Henseler, M., Sandhoff, K., Nakayasu, H. and Suzuki, K. (1996). Targeted disruption of the mouse sphingolipid activator protein gene: a complex phenotype, including severe leukodystrophy and wide-spread storage of multiple sphingolipids. Hum. Mol. Genet. 5, 711–725.

32. Oya, Y., Nakayasu, H., Fujita, N., Suzuki, K. and Suzuki, K. (1998). Pathological study of mice with total deficiency of sphingolipid activator proteins (SAP knockout mice). Acta. Neuropathol. 96, 29–40.

33. Yuan, C., Rao, R.P., Jesmin, N., Bamba, T., Nagashima, K., Pascual, A., Preat, T., Fukusaki, E., Acharya, U. and Acharya, J.K. (2011). CDase is a pan-ceramidase in Drosophila. Mol. Biol. Cell 22(1), 33–43.

34. Heisenberg, M. (1971). Separation of receptor and lamina potentials in the electroretinogram of normal and mutant Drosophila. J. Exp. Biol. 55, 85–100.

35. Coombe, P.E. (1986). The large monopolar cells L1 and L2 are responsible for the ERG transients in Drosophila. J. Comp. Physiol. 159, 655–666.

36. Scott, K. and Zuker, C. (1997). Lights out: deactivation of the phototransduction cascade. Trends. Biochem. Sci. 22, 350–354.

37. Rajaram, S., Scott, R.L. and Nash, H.A. (2005). Retrograde signaling from the brain to the retina modulates the termination of the light response in Drosophila. Proc. Natl. Acad. Sci. U S A. 102, 17840–17845.

38. Ginzburg, L., Kacher, Y. and Futerman, A.H. (2004). The pathogenesis of glycosphingolipid storage disorders. Semin. Cell Dev. Biol. 15, 417–431.

39. Jeyakumar, M., Dwek, R.A., Butters, T.D. and Platt, F.M. (2005). Storage solutions: treating lysosomal disorders of the brain. Nat. Rev. Neurosci. 6, 713–725.

40. Lloyd-Evans, E., Morgan, A.J., He, X., Smith, D.A., Elliot-Smith, E., Sillence, D.J., Churchill, G.C., Schuchman, E.H., Galione, A. and Platt, F.M. (2008). Niemann-Pick disease type C1 is a sphingosine storage disease that causes deregulation of lysosomal calcium. Nat. Med. 14, 1247–1255.

41. Lloyd-Evans, E., Pelled, D., Riebeling, C., Bodennec, J., de-Morgan, A., Waller, H., Schiffmann, R. and Futerman, A.H. (2003). Glucosylceramide and glucosylsphingosine modulate calcium mobilization from brain microsomes via different mechanisms. J. Biol. Chem. 278, 23594–23599.

42. Lloyd-Evans, E., Pelled, D., Riebeling, C. and Futerman, A.H. (2003). Lyso-glycosphingolipids mobilize calcium from brain microsomes via multiple mechanisms. Biochem. J. 375, 561–565.

43. Pelled, D., Lloyd-Evans, E., Riebeling, C., Jeyakumar, M., Platt, F.M. and Futerman, A.H. (2003). Inhibition of calcium uptake via the sarco/endoplasmic reticulum Ca2+-ATPase in a mouse model of Sandhoff disease and prevention by treatment with N-butyldeoxynojirimycin. J. Biol. Chem. 278, 29496–29501.

44. Pelled, D., Trajkovic-Bodennec, S., Lloyd-Evans, E., Sidransky, E., Schiffmann, R. and Futerman, A.H. (2005). Enhanced calcium release in the acute neuronopathic form of Gaucher disease. Neurobiol. Dis. 18, 83–88.

45. Wang, T., Xu, H., Oberwinkler, J., Gu, Y., Hardie, R.C. and Montell, C. (2005). Light activation, adaptation, and cell survival functions of the Na+/Ca2+ exchanger CalX. Neuron 45(3), 367–378

46. Hazkani-Covo, E., Altman, N., Horowitz, M. and Graur, D. (2002). The evolutionary history of prosaposin: two successive tandem-duplication events gave rise to the four saposin domains in vertebrates. J. Mol. Evol. 54, 30–34.

47. Freeman, M.R., Delrow, J., Kim, J., Johnson, E. and Doe, C.Q. (2003). Unwrapping glial biology: Gcm target genes regulating glial development, diversification, and function. Neuron 38, 567–580.

48. Hineno, T., Sano, A., Kondoh, K., Ueno, S., Kakimoto, Y. and Yoshida, K. (1991). Secretion of sphingolipid hydrolase activator precursor, prosaposin. Biochem. Biophys. Res. Commun. 176, 668–674.

49. Kondoh, K., Hineno, T., Sano, A. and Kakimoto, Y. (1991). Isolation and characterization of prosaposin from human milk. Biochem. Biophys. Res. Commun. 181, 286–292.

50. Hiraiwa, M., O'Brien, J.S., Kishimoto, Y., Galdzicka, M., Fluharty, A.L., Ginns, E.I. and Martin, B.M. (1993). Isolation, characterization, and proteolysis of human prosaposin, the precursor of saposins (sphingolipid activator proteins). Arch. Biochem. Biophys. 304, 110–116.

51. Patton, S., Carson, G.S., Hiraiwa, M., O'Brien, J.S. and Sano, A. (1997). Prosaposin, a neurotrophic factor: presence and properties in milk. J. Dairy Sci. 80, 264–272.

52. Sylvester, S.R., Morales, C., Oko, R. and Griswold, M.D. (1989). Sulfated glycoprotein-1 (saposin precursor) in the reproductive tract of the male rat. Biol. Reprod. 41, 941–948.

53. Morales, C.R., Zhao, Q., El-Alfy, M. and Suzuki, K. (2000). Targeted disruption of the mouse prosaposin gene affects the development of the prostate gland and other male reproductive organs. J. Androl. 21, 765–775.

54. Sun, Y., Witte, D.P., Zamzow, M., Ran, H., Quinn, B., Matsuda, J. and Grabowski, G.A. (2007). Combined saposin C and D deficiencies in mice lead to a neuronopathic phenotype, glucosylceramide and alpha-hydroxy ceramide accumulation, and altered prosaposin trafficking. Hum. Mol. Genet. 16, 957–971.

55. Levade, T., Moser, H.W., Fensom, A.H., Harzer, K., Moser, A.B. and Salvayre, R. (1995). Neurodegenerative course in ceramidase deficiency (Farber disease) correlates with the residual lysosomal ceramide turnover in cultured living patient cells. J. Neurol. Sci. 134, 108–114.

56. Matsuda, J., Kido, M., Tadano-Aritomi, K., Ishizuka, I., Tominaga, K., Toida, K., Takeda, E., Suzuki, K. and Kuroda, Y. (2004). Mutation in saposin D domain of sphingolipid activator protein gene causes urinary system defects and cerebellar Purkinje cell degeneration with accumulation of hydroxy fatty acid-containing ceramide in mouse. Hum. Mol. Genet. 13, 2709–2723.

57. Fewou, S.N., Bussow, H., Schaeren-Wiemers, N., Vanier, M.T., Macklin, W.B., Gieselmann, V. and Eckhardt, M. (2005). Reversal of non-hydroxy:alpha-hydroxy galactosylceramide ratio and unstable myelin in transgenic mice overexpressing UDP-galactose:ceramide galactosyltransferase. J. Neurochem. 94, 469–481.

58. Sun, Y., Witte, D.P., Ran, H., Zamzow, M., Barnes, S., Cheng, H., Han, X., Williams, M.T., Skelton, M.R., Vorhees, C.V. et al. (2008). Neurological deficits and glycosphingolipid accumulation in saposin B deficient mice. Hum. Mol. Genet. 17, 2345–2356.

59. Matsuda, J., Vanier, M.T., Saito, Y., Tohyama, J., Suzuki, K. and Suzuki, K. (2001). A mutation in the saposin A domain of the sphingolipid activator protein (prosaposin) gene results in a late-onset, chronic form of globoid cell leukodystrophy in the mouse. Hum. Mol. Genet. 10, 1191–1199.

60. Sun, Y., Ran, H., Zamzow, M., Kitatani, K., Skelton, M.R., Williams, M.T., Vorhees, C.V., Witte, D.P., Hannun, Y.A. and Grabowski, G.A. (2010). Specific saposin C deficiency: CNS impairment and acid beta-glucosidase effects in the mouse. Hum. Mol. Genet. 19, 634–647.

61. Sun, Y., Liou, B., Ran, H., Skelton, M.R., Williams, M.T., Vorhees, C.V., Kitatani, K., Hannun, Y.A., Witte, D.P., Xu, Y.H. et al. (2010). Neuronopathic Gaucher disease in the mouse: viable combined selective saposin C deficiency and mutant glucocerebrosidase (V394L) mice with glucosylsphingosine and glucosylceramide accumulation and progressive neurological deficits. Hum. Mol. Genet. 19, 1088–1097.

62. Min, K.T. and Benzer, S. (1997). Spongecake and eggroll: two hereditary diseases in Drosophila resemble patterns of human brain degeneration. Curr. Biol. 7, 885–888.

63. Nakano, Y., Fujitani, K., Kurihara, J., Ragan, J., Usui-Aoki, K., Shimoda, L., Lukacsovich, T., Suzuki, K., Sezaki, M., Sano, Y. et al. (2001). Mutations in the novel membrane protein spinster interfere with programmed cell death and cause neural degeneration in Drosophila melanogaster. Mol. Cell Biol. 21, 3775–3788.

64. Hickey, A.J., Chotkowski, H.L., Singh, N., Ault, J.G., Korey, C.A., MacDonald, M.E. and Glaser, R.L. (2006). Palmitoyl-protein thioesterase 1 deficiency in Drosophila melanogaster causes accumulation of abnormal storage material and reduced life span. Genetics 172, 2379–2390.

65. Dawkins, J.L., Hulme, D.J., Brahmbhatt, S.B., Auer-Grumbach, M. and Nicholson, G.A. (2001). Mutations in SPTLC1, encoding serine palmitoyltransferase, long chain base subunit-1, cause hereditary sensory neuropathy type I. Nat. Genet. 27, 309–312.

66. Myllykangas, L., Tyynela, J., Page-McCaw, A., Rubin, G.M., Haltia, M.J. and Feany, M.B. (2005). Cathepsin D-deficient Drosophila recapitulate the key features of neuronal ceroid lipofuscinoses. Neurobiol. Dis. 19, 194–199.

67. Tuxworth, R.I., Vivancos, V., O'Hare, M.B. and Tear, G. (2009). Interactions between the juvenile Batten disease gene, CLN3, and the Notch and JNK signalling pathways. Hum. Mol. Genet. 18, 667–678.

68. Yonamine, I., Bamba, T., Nirala, N.K., Jesmin, N., Kosakowska-Cholody, T., Nagashima, K., Fukusaki, E., Acharya, J.K. and Acharya, U. (2011). Sphingosine kinases and their metabolites modulated endolysosomal trafficking in photoreceptors. J. Cell Biol. 192(4), 557–567.

69. Matyash, V., Liebisch, G., Kurzchalia, T.V., Shevchenko, A. and Schwudke, D. (2008). Lipid extraction by methyl-tert-butyl ether for high-throughput lipidomics. J. Lipid Res. 49(5), 1137–1146.

70. Schwudke, D., Schuhmann, K., Herzog, R., Bornstein, S.R. and Shevchenko, A. (2011). Shotgun lipidomics on high resolution mass spectrometers. Cold Spring Harb. Perspect. Biol. 3(9),a004614

71. Herzog, R., Schwudke, D., Schuhmann, K., Sampaio, J.L., Bornstein, S.R., Schroeder, M., Shevchenko, A. (2011). A novel informatics concept for high-throughput shotgun lipidomics based on the molecular fragmentation query language. Genome Biol. 12(1), R8

72. Hindle, S., Afsari, F., Stark, M., Middleton, C.A., Evans, G.J., Sweeney, S.T. and Elliott, C.J. (2013). Dopaminergic expression of the Parkinsonian gene LRRK2-G2019S leads to non-autonomous visual neurodegeneration, accelerated by increased neuronal demands for energy. Hum. Mol. Genet. 22(11), 2129–2140.

73. Zafeiriou DI, Triantafyllou P, Gombakis NP, Vargiami E, Tsantali C and Michelakaki E. (2003) Niemann-Pick type C disease associated with peripheral neuropathy. Pediatr. Neurol. 29(3), 242–244

